# ARCADE disentangles cell abundance from continuous transcriptional state in spatial transcriptomics

**DOI:** 10.64898/2026.10.28.762703

**Authors:** Qiang Su, Yi Long, Jing Jiang, Chen Yang, Qizhou Lian

## Abstract

Spatial transcriptomics is widely used to infer both cell-type abundance and within-type transcriptional variation, but these signals are difficult to separate in mixed spot-level measurements. Here, we present ARCADE, a computational framework that addresses two major sources of uncertainty in spatial analysis: unstable single-cell references and entanglement between abundance and cell state. ARCADE first builds a robust reference through hyperparameter optimization and consensus filtering, then performs spatial inference in two stages by estimating cell-type composition before modeling continuous intratype transcriptional programs around this compositional anchor. Across simulated and semi-synthetic benchmarks, ARCADE improves deconvolution accuracy and yields more stable state inference than existing approaches. In matched human dorsolateral prefrontal cortex datasets with spatial proteogenomic readouts, ARCADE distinguishes localized cell enrichment from astrocyte and microglial activation programs. These results show that decoupled modeling can improve the interpretability of spatial transcriptomic signals in complex tissues.

## Introduction

Spatial transcriptomics (ST) has enabled transcriptome-wide measurement of tissue organization while preserving spatial context, providing a powerful framework for studying development, physiology, and disease in situ.^1-3^ However, for widely used sequencing-based ST platforms, each spatial capture location typically contains transcripts from multiple cells rather than a single resolved cell.^4-6^ As a result, computational analysis must infer the cellular composition of each spot from mixed expression profiles before higher-order spatial programs can be interpreted. This creates two related but distinct analytical objectives. The first is spatial deconvolution: estimating the proportions of discrete cell types within each spot. The second is state-resolved spatial modeling: characterizing continuous transcriptional variation within a given cell type across locations, such as local immune activation, stress responses, metabolic adaptation, or differentiation gradients.^7-9^ While the first objective is now addressed by a mature class of methods, the second remains substantially more challenging, particularly when both abundance and intratype state are inferred from the same mixed measurements.

Existing methods can be broadly organized into three categories. A first class includes deconvolution approaches such as RCTD^10^, Cell2location^11^, Stereoscope^12^, SPOTlight^13^, and Seurat-based transfer pipelines.^14^ These methods are effective for estimating broad cellular composition, but they generally assume relatively fixed reference profiles and therefore do not explicitly model continuous state variation within cell types. A second class includes spatial mapping methods such as Tangram^15^ and CellTrek^16^ which transfer cells or transcriptional profiles from single-cell references into spatial coordinates. These approaches are useful for reference-guided charting, but their inferences remain constrained by the states represented in the input single-cell dataset and are not designed to learn previously unobserved spatially induced expression programs. A third class includes joint generative models such as DestVI^17^ and Starfysh^18^, which attempt to infer both composition and continuous cell states from spatial data. These methods represent an important conceptual advance, but they also face a difficult identifiability problem: the same local expression shift can often be explained either by a change in cell-type abundance or by a change in the transcriptional state of cells already present in the spot. More generally, disentangled latent representations in variational models are not guaranteed to be identifiable without additional assumptions or inductive constraints.^19, 20^

This ambiguity is especially important when expression changes are concentrated in canonical marker genes. For example, an increase in a lineage marker within a spatial region may reflect accumulation of that cell type, activation of a subset of cells within that lineage, or both. When abundance and state are optimized jointly without a sufficiently strong compositional constraint, the model may distribute signal between these components in unstable or biologically implausible ways. In this study, we refer to this failure mode as state leakage: inferred continuous state variation spuriously tracks inferred abundance, or vice versa, because both are recruited to explain the same transcriptional deviation. Importantly, some correlation between abundance and state can be biologically genuine, so the challenge is not to eliminate all dependence, but to reduce model-induced entanglement and improve the interpretability of each component. A second and often underappreciated challenge lies upstream of spatial inference itself: the construction of a reliable single-cell reference. Many deconvolution and state-aware spatial frameworks depend directly on cell-type definitions or expression programs derived from scRNA-seq, while other approaches rely on marker genes or prior annotations whose utility is shaped by the quality and transferability of upstream single-cell assignments.^14, 17, 21^ In practice, these assignments can vary substantially with choices made during preprocessing and clustering, including feature selection depth, dimensionality reduction, neighborhood graph construction, and resolution parameters.^14, 21-26^ Importantly, this upstream instability should be distinguished from biologically meaningful continuous state variation. Cells occupying activated, transitional, or other noncanonical transcriptional states are not inherently problematic for spatial modeling if they can still be assigned with confidence to a defined cell type or lineage context. The more relevant concern is annotation instability: cells whose identity remains inconsistent across complementary assignment views, such as cell-level prediction, cluster-level topology, and iterative refinement. If these unstable assignments are incorporated directly into the reference, they can blur the compositional backbone used for downstream spatial inference and exacerbate abundance-state entanglement. Because these sources of variation propagate into downstream spatial models, instability in the reference can amplify uncertainty in both deconvolution and inferred cell states.

We therefore sought to develop a framework that addresses both problems in a coordinated manner: improving the robustness of the upstream reference while explicitly separating discrete composition from continuous state during spatial inference. Here we introduce ARCADE (Auto-optimized Reference Consensus And Decoupling Engine), a cross-ecosystem computational pipeline for spatial transcriptomic analysis. In the reference-construction stage, ARCADE combines Bayesian hyperparameter optimization with consensus filtering to identify preprocessing and clustering settings that yield a more stable identity backbone. In this context, consensus filtering is used not to remove all transcriptionally unusual cells, but to reduce cells with persistently inconsistent identity assignments across complementary annotation views. Depending on the dataset and annotation setting, this can be implemented as bi-consensus or tri-consensus filtering. It then applies a two-stage spatial model in which cell-type proportions are first estimated using a constrained autoencoder to provide a compositional anchor, after which a variational model infers residual intratype transcriptional programs under regularization around that anchor. By separating these steps, ARCADE is designed to reduce the non-identifiability that can arise when abundance and state are inferred simultaneously from the same spot-level measurements. Using controlled simulations, semi-synthetic benchmarks, and matched human dorsolateral prefrontal cortex (DLPFC) datasets with orthogonal spatial proteogenomic measurements, we show that ARCADE improves compositional accuracy and yields more stable, biologically interpretable spatial state inference. In particular, ARCADE supports the distinction between localized cellular enrichment and localized transcriptional activation in astrocyte and microglial programs, illustrating how decoupled modeling can refine interpretation of spatial transcriptomic signals in complex tissues.

## Results

### Joint inference induces abundance-state entanglement in controlled settings

Reliable inference of spatial cell-type abundance and continuous intratype transcriptional state depends in part on the quality of the single-cell reference used to guide analysis. In practice, however, standard scRNA-seq preprocessing workflows remain sensitive to several interdependent hyperparameters, including the number of highly variable genes (HVGs), the dimensionality of principal component analysis, the construction of the k-nearest neighbor graph, and clustering resolution. These choices are often made sequentially using heuristic defaults, even though they jointly influence cluster structure, annotation stability, and the distinction between well-resolved and ambiguous cell populations. As a result, uncertainty in the reference can propagate into downstream spatial inference, affecting both deconvolution and the interpretation of inferred cell states.

To address this upstream source of variability while supporting abundance-state decoupling downstream, we developed ARCADE as a unified framework that combines reference optimization with two-stage spatial inference (Figure 1A). The workflow begins with Gene Universe Harmonization (GUH), followed by Bayesian optimization (BO) and Consensus Filtering (CF) to construct a more stable scRNA-seq reference. This optimized reference is then used in a decoupled spatial model in which cell-type composition is first estimated to define a compositional anchor, after which continuous within-type transcriptional variation is inferred under regularization around that anchor. ARCADE first applies GUH to restrict the feature space to genes detected in both the single-cell and spatial datasets (*G*_Common_ = *G*_*sc*_*rn*_*A*_ ∩ *G*_st_). This step reduces reliance on modality-specific features that cannot contribute to downstream spatial reconstruction and helps align the reference with the transcriptomic space that is observable in the spatial data. ARCADE then performs joint hyperparameter search using a multi-strategy Bayesian optimization engine that explores combinations of HVG number, principal component dimension, k-nearest neighbor connectivity, and clustering resolution (Figure 1A and 1B). Rather than tuning these parameters independently, ARCADE evaluates each candidate configuration as a complete preprocessing-and-clustering solution.

**Figure 1.**
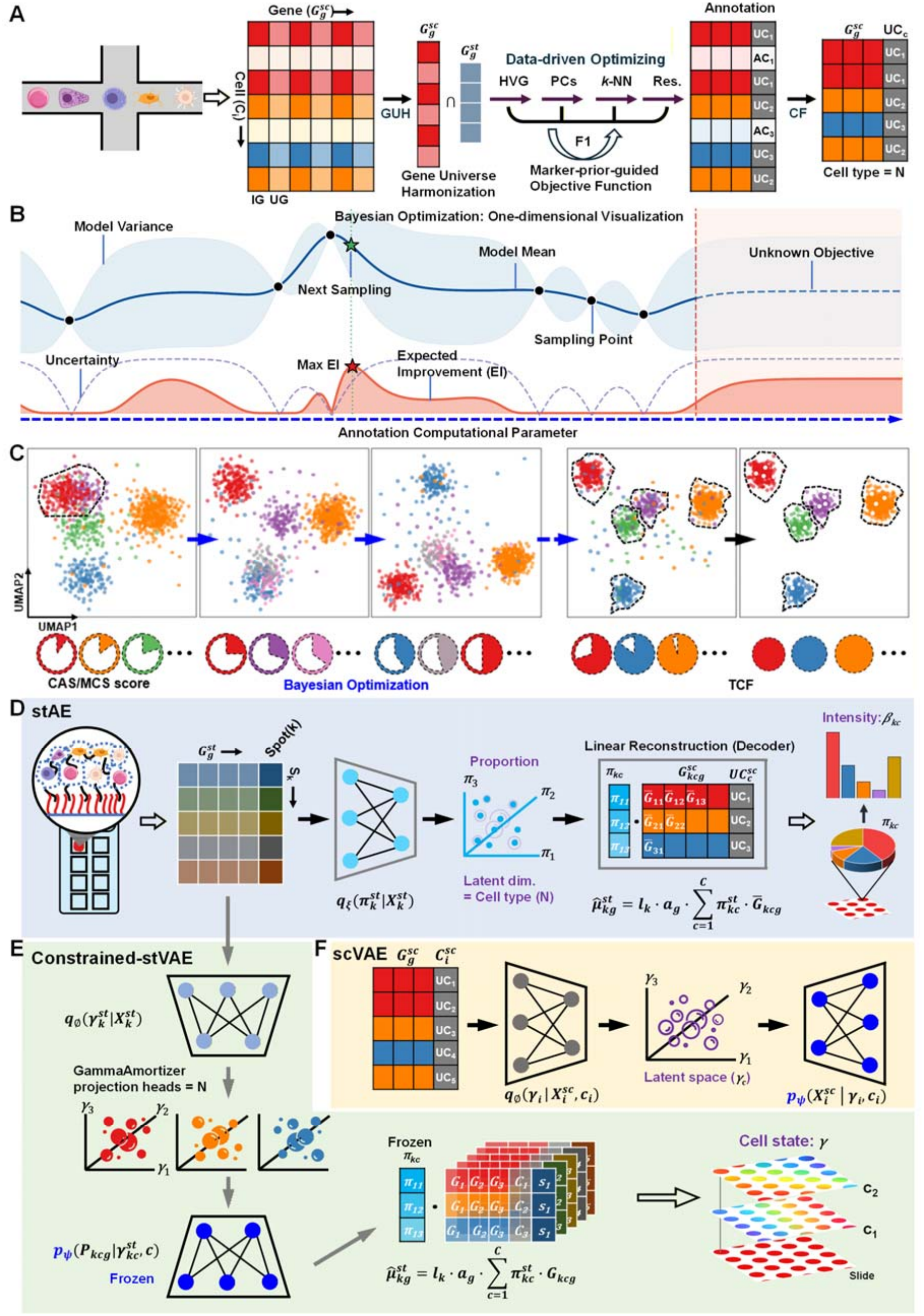
The ARCADE framework for Bayesian-optimized reference generation and decoupled spatial state inference. **A**) Schematic of the ARCADE workflow. Quality-controlled scRNA-seq matrices first undergo Gene Universe Harmonization (GUH), restricting the feature space to genes shared with the spatial transcriptomics (ST) query (*G*_common_ *= G*_scRNA_ *∩ G*_*ST*_) . The harmonized matrices then enter a Bayesian optimization (BO) procedure that jointly tunes interdependent preprocessing and clustering hyperparameters, including the number of highly variable genes, principal components, k-nearest neighbor graph settings, and clustering resolution. Candidate solutions are evaluated using a marker-informed objective together with data-driven clustering criteria. The resulting reference is further refined by Consensus Filtering (CF), implemented as either bi-consensus or tri-consensus filtering depending on the annotation layers used. Cells with consistent assignments across the configured annotation views are retained to generate the final filtered reference, which is subsequently used for decoupled spatial inference. **B**) Mechanics of the Bayesian hyperparameter search. The upper plot illustrates the Gaussian process (GP) surrogate model, showing the posterior mean *(µ)* and uncertainty (σ) estimated from observed evaluations. The lower plot shows the expected improvement (EI) acquisition function together with uncertainty magnitude. By selecting the next parameter configuration that maximizes EI, ARCADE balances exploitation of promising regions with exploration of uncertain regions of the hyperparameter space. **C**) Illustration of the optimization trajectory across clustering states. The BO engine searches for parameter configurations that improve a composite objective integrating the Cluster Annotation Score (CAS), Marker Consistency Score (MCS), and silhouette score. This procedure tends to disfavor both under-clustered solutions that merge biologically distinct populations and over-clustered solutions that fragment continuous variation or technical noise. **D**) The spatial autoencoder (stAE) used for deconvolution and compositional anchoring. A neural encoder compresses the spatial gene expression matrix (*X*^*st*^) to infer spot-level cell-type proportions (*π*^*st*^). which are constrained to a sum-to-one simplex. The decoder reconstructs expected expression 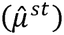 through a linear mixture model combining the optimized scRNA-seq reference profiles, spot-specific library sizes. and platform-adaptive gene correction factors (α). At this stage. reference populations are treated as fixed aggregates, so the model focuses on composition without explicitly inferring continuous cell state. **E**) The spatial variational autoencoder (stVAE) for decoupled cell-state inference. The stAE-derived proportions *(π*^*st*^) are incorporated as a soft-constraint prior during spatial state inference. A variational GammaAmortizer maps each spatial spot to latent cell-state variables (*γ*^*st*^) through a panel of *N* cell-type-specific projection heads. where *N* denotes the total number of reference cell types. Each head outputs the mean and variance parameters for the latent state associated with its corresponding cell type. During reconstruction, the transferred cell-type-conditioned decoder p_θ_ is applied in parallel across cell types. and the decoded outputs are combined according to *(π*^*st*^) . This architecture encourages expression variation not explained by composition to be represented in latent state coordinates. **F**) The single-cell variational autoencoder (scVAE) used for reference manifold learning. A conditional VAE is trained on the filtered single-cell reference *(X*^*sc*^) together with cell-type annotations to learn a latent representation of within-cell-type transcriptional variation (*γ*^*sc*^). The encoder uses the same variational parameterization as the spatial model, and the resulting cell-type-conditioned decoder p_θ_ is subsequently transferred to the stVAE and held fixed during spatial state inference. providing a shared reference-informed generative basis across modalities.

To assess candidate solutions, ARCADE is able to use a modular objective function that combines biological priors with data-driven structure. When canonical marker information is available, the search is guided in part by a Marker Prior Score (MPS), which measures concordance between cluster-level differentially expressed genes and expected marker sets using an F1-based criterion. In this formulation, precision rewards clusters whose defining genes align with expected lineage markers, whereas recall rewards recovery of the expected marker repertoire for each cell type. Because marker priors may be incomplete or context-dependent, ARCADE combines MPS with a data-driven composite score that captures annotation and cluster coherence. This composite score includes the Cluster Annotation Score (CAS), which quantifies agreement between cell-level predictions and cluster-level consensus labels; the Marker Consistency Score (MCS), which evaluates the prevalence and coherence of de novo marker expression within clusters; and the silhouette score, which measures geometric separation in latent space (Figure 1C). Together, these terms prioritize candidate references that are both biologically interpretable and structurally coherent. To improve exploration of the hyperparameter landscape, ARCADE executes multiple BO strategies in parallel, emphasizing exploratory, exploitative, and balanced search behaviors. Information from these search trajectories is integrated through a shared Gaussian process (GP) surrogate model, and candidate updates are selected using an expected improvement (EI) acquisition function (Figure 1B). This design allows the optimization procedure to balance exploration of uncertain regions against exploitation of promising parameter configurations. In practice, the search tends to disfavor both under-resolved solutions, in which biologically distinct populations are merged, and highly fragmented solutions, in which technical noise or continuous transitional variation is split into excessive clusters (Figure 1C).

Because hyperparameter optimization alone does not fully resolve uncertainty arising from unstable cell-type assignments, ARCADE next applies Consensus Filtering (CF) to refine the reference. In this framework, we distinguish continuous transcriptional state variation from annotation ambiguity. Cells exhibiting transitional, activated, or otherwise noncanonical programs are not considered ambiguous if they can still be assigned confidently to a defined cell type or lineage context. By contrast, CF targets cells whose annotations remain inconsistent across complementary assignment views, including cell-level prediction, topology-based cluster consensus, and iterative refinement procedures. Depending on the dataset and annotation setting, this can be implemented as bi-consensus or tri-consensus filtering. The aim is not to produce a universally “pure” reference, but to reduce instability in the identity backbone while preserving biologically meaningful intratype variation for downstream spatial modeling. ARCADE then uses this optimized reference in a decoupled two-stage spatial architecture designed to reduce the ambiguity that arises when cell abundance and continuous state are inferred simultaneously from the same spot-level measurements. As discussed above, local expression changes can often be explained either by altered cell-type proportions or by altered transcriptional state within a cell type. When both components are optimized jointly without a sufficiently strong compositional constraint, signal may be distributed between them in unstable ways. We refer to this model-induced failure mode as state leakage.

In the first stage, ARCADE estimates spatial composition using a spatial autoencoder (stAE) that provides a compositional anchor for each spot (Figure 1D). The encoder maps spot-level spatial expression (*X*^*st*^) into a low-dimensional latent representation whose dimensionality matches the number of reference cell types, yielding estimated spot-level proportions (*π*^*st*^). A sum-to-one simplex constraint encourages these latent variables to behave as cell-type abundances. The decoder reconstructs expected spatial expression 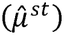 through a linear mixture model that combines the optimized scRNA-seq reference profiles with spot-specific library sizes and platform-adaptive gene correction factors (*α*). At this stage, reference populations are treated as fixed aggregates and continuous state variation is not explicitly modeled, allowing the stAE to focus on recovering broad compositional structure without allocating local residual variation to cell-state effects prematurely. In the second stage, ARCADE models continuous transcriptional variation using a spatial variational autoencoder (stVAE) while regularizing around the compositional anchor estimated in stage one (Figure 1E). To define the space of plausible within-type transcriptional programs, ARCADE first trains a conditional single-cell VAE (scVAE) on the filtered reference, using single-cell expression (*X*^*sc*^) together with cell-type annotations to learn latent state variables (*γ*^*sc*^) conditioned on cell identity (Figure 1F). The resulting cell-type-conditioned decoder *p*_*θ*_ is then transferred to the spatial setting and held fixed during spatial state inference, providing a shared generative basis across modalities. During spatial inference, the stVAE uses the stage-one abundance estimates (*π*^*st*^) as a soft-constraint prior rather than a hard constraint, so that abundance estimates may adjust modestly when strongly supported by the data but are discouraged from drifting substantially away from the compositional anchor. A dedicated variational module, implemented here as a GammaAmortizer, maps each spatial spot to latent cell-state variables (*γ*^*st*^). This amortizer consists of a panel of *N* cell-type-specific projection heads, where *N* is the number of reference cell types, and each head generates the mean and variance parameters for the latent coordinate associated with its corresponding cell type. During reconstruction, the transferred decoder *p*_*θ*_ is invoked in parallel across all cell types, and the decoded outputs are linearly combined according to the abundance prior (*π*^*st*^). Under this architecture, residual expression variation not adequately explained by composition is encouraged to be represented in the latent state coordinates, supporting interpretation in terms of continuous in situ transcriptional plasticity rather than arbitrary reshaping of cell-type abundance.

In addition to abundance-state decoupling, this generative framework provides useful downstream summaries for spatial analysis. ARCADE estimates cell-type-specific imputed expression at each spatial location by combining inferred composition with the state-conditioned decoder, enabling inspection of gene programs within individual cell types even when they represent a small fraction of a spot. ARCADE also produces a denoised tissue-level reconstruction that integrates inferred abundance, latent state, and platform-adaptive correction terms. We view these outputs as model-based reconstructions intended to improve interpretability of sparse spatial measurements rather than as direct replacements for observed data. Together, these components position ARCADE as a unified framework in which robust reference construction supports decoupled inference of cell abundance and continuous transcriptional state in spatial transcriptomics.

### Bayesian reference optimization stabilizes the compositional backbone for spatial inference

Because ARCADE uses a single-cell reference to anchor downstream spatial inference, we first examined whether common preprocessing choices alter reference annotation in ways that could propagate into deconvolution and state estimation. We systematically titrated the number of highly variable genes (HVGs) in public 10x Genomics benchmark datasets, including 5k Mouse Brain CNIK 3’ v3.1 and 10k Human PBMC v3, and compared annotations to the full-transcriptome baseline. In both datasets, restricting the feature space substantially reduced cell-level concordance, although the pattern of instability differed by tissue context. In mouse brain, exact-match annotation accuracy declined from 98.73% at 20,000 genes to 13.67% at 500 genes, with progressive collapse of fine regional and subtype distinctions (Figure S1-3). In PBMC, exact-match accuracy similarly decreased from 90.47% to 16.69%, but the dominant effect was redistribution among closely related immune states rather than broad lineage failure (Figure S4 and 5). Barcode-level tracking further showed that annotation stabilization occurred more slowly in brain than in PBMC, with only 38.3% of brain cells but 73.1% of PBMC cells retaining their final identity by 3,000 HVGs (Figure S3). Repeating the analysis with Seurat’s anchor-based annotation pipeline yielded the same general conclusion: broad category counts appeared relatively stable, but cell-level assignments remained sensitive to feature selection (Figure S6).

These observations motivated the reference optimization component of ARCADE. In a simulated scRNA-seq dataset comprising 15,000 cells across 10 transcriptionally distinct groups Generated by combining Splatter^27^ and Polyester^28^, we found that clustering quality depended nonlinearly on the joint choice of HVG number, principal component depth, neighborhood size, and clustering resolution (Figure S7). Parameter interactions produced response surfaces that were not well captured by sequential one-parameter tuning, supporting the use of Bayesian optimization to search the coupled hyperparameter space. ARCADE implements this step using a Gaussian process-guided strategy that balances exploration and exploitation to identify reference configurations with improved topological agreement. We next applied this framework to an in-house mouse striatum scRNA-seq reference containing 15,282 cells and 187 high-granularity initial cell-level annotations. Across a 90-trial Bayesian search, candidate parameter settings yielded markedly different reference topologies, spanning 10 to 17 consensus clusters, while the composite optimization objective increased from approximately 71.7% to >80.3% (Figure S8; Figure S9a). Representative solutions illustrated several recurring failure modes, including over-merging at low resolution and unstable fragmentation under sparse-feature, high-resolution settings. Together, these analyses indicate that heuristic preprocessing can substantially reshape the compositional backbone used for spatial inference, and that Bayesian optimization provides a more stable reference construction procedure for the downstream decoupling framework.

One early configuration characterized by high connectivity and low clustering resolution (*n*_HVG_ = 4,079, *n*_PC_ = *69, k* = 45, resolution = 0.2383) compressed the dataset into only 7 consensus clusters (Figure S9b; Figure S10), merging neuronal populations that were better separated under other settings. A different intermediate configuration with relatively restricted latent dimensionality (*n*_HvG_ = 7,073, *n*_PC_ = 10, *k* = 29, resolution = 1.1148) produced 11 consensus clusters and showed limited ability to resolve closely related glutamatergic populations (Figure 2A; Figure S11). For example, the “151 TH Prkcd Grin2c Glut” cluster contained 5,161 cells, of which 2,953 matched the individual predictions ( CAS = 57.2% ), while the “150 CM-IAD-CL-PCN Sema5b Glut” and “154 PF Fzd5 Glut” clusters showed lower assignment concordance (39.8%and 28.5%, respectively). Conversely, a high-partition, low-feature configuration (*n*_HVG_ = 800,7*n*_PC_ = 100, *k =* 10, resolution = 1.8208) yielded 16 consensus clusters but remained unstable among inhibitory populations, including a “101 ZI Pax6 Gaba” cluster with a CAS of 11.0% (Figure S9c; Figure S12). Notably, in several of these cases, the corresponding Marker Consistency Score (MCS) remained high despite lower Consensus Assignment Score (CAS) values, indicating that marker expression alone may not fully capture errors introduced by suboptimal topology. Under the best-performing parameter configuration identified by the composite objective (*n*_HVG_ = 5,769,*n*_PC_ = 50, *k* = 21, resolution = 1.1322), the reference organized into 14 consensus clusters with improved agreement between local topology and cell-level annotation (Figure 2B; Figure S13). In this setting, previously compressed glutamatergic populations became better separated, and the “154 PF Fzd5 Glut” cluster improved from a CAS of 28.5% under the suboptimal configuration to 92.6%, while maintaining high marker consistency (MCS = 97.3%). The “327 Oligo NN” cluster remained stable across settings, indicating that some lineages were less sensitive to hyperparameter choice than others. These comparisons suggest that optimization is particularly important for resolving populations whose boundaries are not readily preserved under default or heuristic settings.

**Figure 2.**
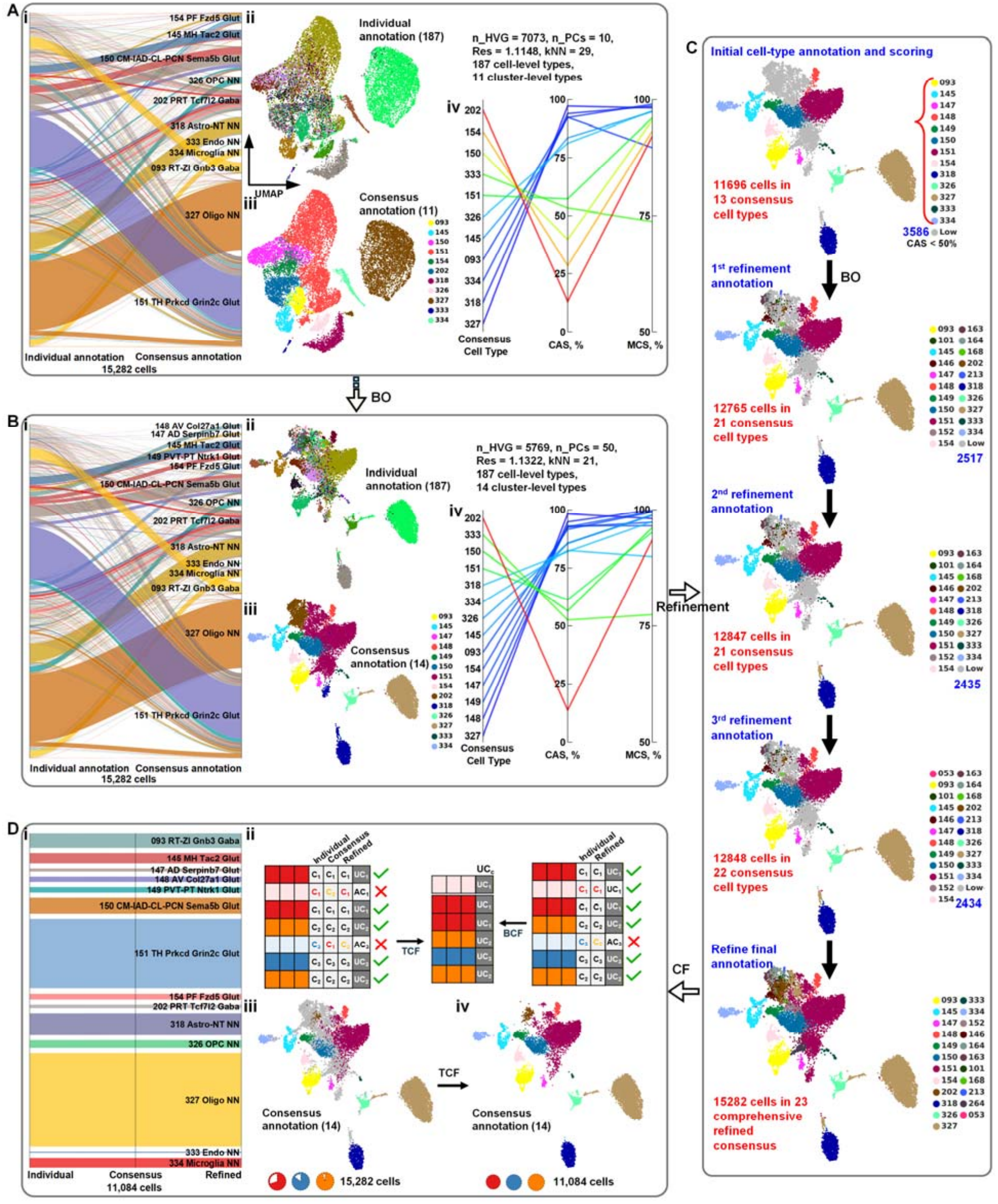
Bayesian hyperparameter optimization, topology-aware annotation assessment, and consensus filtering for reference generation. **A)** Example of a suboptimal Bayesian optimization trial with limited latent dimensionality (*n*_HvG_ = 7,073, *n*_PC:_ = 10,*k* = 29, resolution = 1.11). (i) Alluvial plot showing the mapping of 15,282 cells from 187 high-granularity initial reference annotations to 11 consensus clusters derived from topology-level majority voting. (ii) UMAP projection generated under this parameter setting, colored by the 187 cell-level Individual Predictions. (iii) Corresponding UMAP colored by the 11 Original Consensus cluster labels. (iv) Quantitative evaluation of representative clusters under this configuration. Glial populations such as 327 Oligo NN remain comparatively stable (*n* = 3,985, CAS = 97.2%, MCS = 98.2%), whereas several glutamatergic populations show lower assignment concordance, including 151 TH Prkcd Grin2c Glut (CAS = 57.2%), 150 CM-IAD-CL-PCN Sema5b Glut ( CAS = 39.8%), and 154 PF Fzd5 Glut (CAS = 28.5%). **B)** Best-performing parameter configuration identified by the composite objective (*n*_Hvg_ = 5,769, *n*_PC_ = 50, *k* = 21, resolution = 1.13). (i) Alluvial plot showing the mapping from the 187 initial annotations to 14 consensus clusters under the optimized setting. (ii–iii) UMAP projections of the optimized latent space, illustrating improved separation among previously compressed glutamatergic populations. (iv) Cluster-level evaluation under the optimized configuration. 327 Oligo NN remains stable (u = 3,928, CAS = 98.3%, MCS = 98.2%), while 154 PF Fzd5 Glut shows improved concordance (CAS = 92.6%, MCS = 97.3%). The 202 PRT Tcf7l2 Gaba cluster (*n* = 1,164)illustrates a case of high marker consistency but low assignment concordance (MCS = 93.5%, CAS = 13.7%), motivating local refinement. **C)** Iterative sub-clustering and consensus refinement of locally ambiguous regions. (Top) UMAP of the initial annotation obtained under the optimized setting in B. A total of 11,696 cells are assigned to 13 Original Consensus types exceeding the CAS threshold, while 3,586 cells with lower assignment concordance ( CAS < 50% )are flagged for reanalysis (grey). (Middle) Sequential refinement steps. Flagged cells are iteratively isolated, re-clustered at higher local resolution, and re-evaluated. The first refinement assigns 12,765 cells to 21 types and leaves 2,517 cells unresolved; the second assigns 12,847 cells to 21 types with 2,435 unresolved cells; and the third assigns 12,848 cells to 22 consensus types with 2,434 unresolved cells. (Bottom) Final integrated Refined Annotation across the full dataset (*n =* 15,282), yielding 23 consensus cell types. **D)** Consensus filtering strategies for constructing the final reference backbone. (i) Alluvial plot showing annotation alignment among cells retained after filtering. (ii) Schematic of the filtering rules. Tri-Consensus Filtering (TCF) retains cells for which the Individual Prediction, Original Consensus, and Refined Annotation agree. Bi-Consensus Filtering (BCF) retains cells for which the Individual Prediction agrees with the Refined Annotation. (iii) UMAP of the full pre-filtered dataset (*n* = 15,282 ), colored by TCF retention status. Colored cells indicate the 11,084 high-consistency cells retained by TCF, whereas grey cells indicate cells removed because of conflicting assignments across annotation layers. (iv) UMAP of the TCF-filtered reference, which provides a conservative high-consistency backbone for downstream spatial deconvolution.

We next used the divergence between CAS and MCS to identify regions of the optimized reference that remained locally ambiguous. Rather than interpreting these cells as transitional states, we treated them as cases in which cluster topology and cell-level identity assignment were not yet sufficiently aligned. Clusters with relatively high marker consistency but low assignment concordance (CAS < 50%)were therefore routed to an iterative sub-clustering procedure for local refinement (Figure 2C; Figure S14). At baseline, 11,696 cells mapped to 13 original consensus clusters exceeding the CAS threshold, while 3,586 cells were flagged for reanalysis. The first refinement expanded the high-confidence set to 12,765 cells across 21 types, leaving 2,517 cells unresolved. A second refinement step yielded 12,847 confident cells and 2,435 unresolved cells, and a third iteration reached a stable endpoint with 12,848 confident cells and 2,434 residual unresolved cells. Merging these localized refinements back into the global annotation generated a final Refined Annotation of 23 consensus cell types across the full dataset.

Because optimization and local refinement do not eliminate all cases of unstable identity assignment, ARCADE next applies consensus filtering to define a reference backbone for downstream spatial modeling (Figure 2D; Figure S15). Here we distinguish continuous transcriptional variation within a defined cell type from uncertainty in cell identity. Cells showing activated or transitional programs may still be informative reference examples if they remain consistently assignable to a parent cell type, whereas cells with conflicting assignments across annotation layers are less suitable as reference anchors. To formalize this distinction, we implemented two related filtering strategies. Tri-Consensus Filtering (TCF) retains cells for which the Individual Prediction, Original Consensus, and Refined Annotation all agree (Figure 2D). Applied to the striatal dataset, TCF retained 11,084 cells distributed across 14 lineages, producing a highly consistent core reference. Bi-Consensus Filtering (BCF) relaxes the requirement for agreement with the original consensus and retains cells for which the Individual Prediction matches the Refined Annotation (Figure S15). This strategy preserved 11,899 cells spanning 22 cell types, while excluding 3,382 cells (22.1%) with inconsistent assignments. In practice, TCF provides a conservative reference for applications prioritizing maximal identity stability, whereas BCF retains greater compositional diversity and may better preserve rare or locally obscured populations. Together, these analyses establish the reference optimization and consensus filtering framework used by ARCADE before spatial inference.

### ARCADE accurately recovers abundance and intratype state in biologically coherent semi-synthetic benchmarks

Having established a reference construction procedure for downstream spatial analysis, we next evaluated whether ARCADE could accurately recover both cell-type abundance and continuous intratype transcriptional variation under known ground truth. To do so, we generated large-scale semi-synthetic spatial transcriptomics datasets designed to preserve biologically coherent tissue structure rather than spot-wise random mixtures. Following the spatially aware simulation strategy introduced in DestVI,^17^ we generated smooth spatial domains on a staggered hexagonal lattice matched to the geometry of 10x Genomics Visium. For the primary benchmark, we used a mouse striatum reference derived from the CellTypist-based ARCADE pipeline, comprising 11,048 transcriptomically filtered cells across 22 high-resolution cell types. Cell-type mixing proportions were sampled using Gaussian random fields, producing locally continuous abundance patterns and anatomically plausible transitions across 4,900 capture spots, with 5-10 cells per spot (Figure 3A). Spot-level counts were then simulated using a negative binomial generalized linear model calibrated to empirical sequencing depth and technical variability.

**Figure 3.**
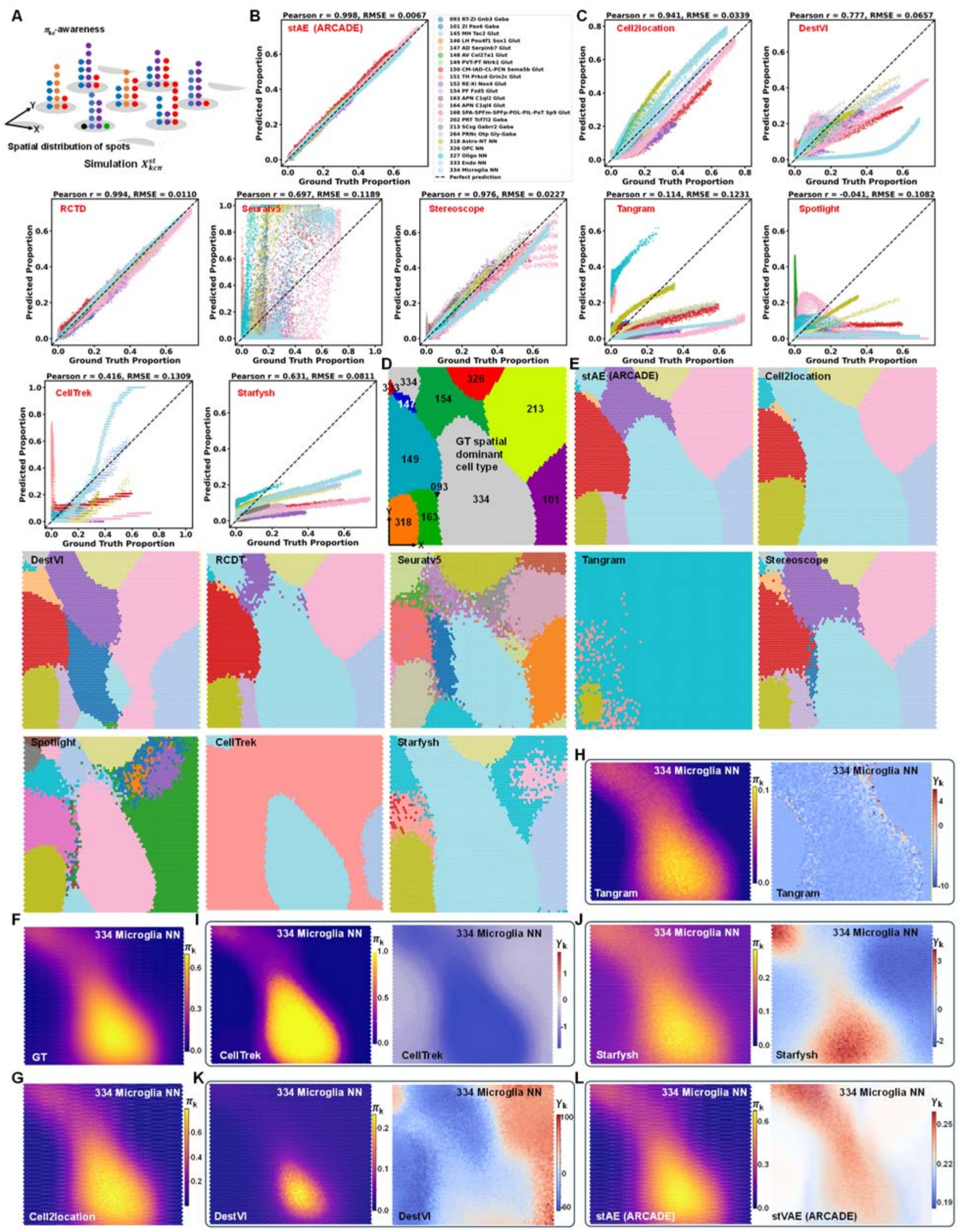
Validation of spatial deconvolution and continuous state inference in biologically coherent semi-synthetic benchmarks. **A)** Construction of the semi-synthetic spatial transcriptomics benchmark (N = 4,900 spots). A mouse striatum single-cell reference generated by the CellTypist-based ARCADE pipeline (11,048 transcriptomically filtered cells across 22 high-resolution cell types) was used to simulate a tissue on a staggered hexagonal lattice matched to 10x Genomics Visium geometry. To preserve spatial continuity, spot-level cell-type proportions (*π*) were sampled from Gaussian random fields, generating anatomically coherent domains with mixtures of 5–10 cells per spot. Gene counts were then simulated using a negative binomial generalized linear model calibrated to empirical sequencing depth and technical variability. **B)** Global deconvolution accuracy of ARCADE across all 107,800 spot-by-cell-type pairs. The scatter plot compares inferred versus true cell-type proportions. ARCADE achieved high global concordance (Pearson r = 0.9978, Spearman ρ = 0.9193, R^2^ = 0.9954, RMSE = 0.0067, MAE = 0.0040). Across the reference, 20 of 22 cell types showed per-cell-type Pearson r > 0.99 (median per-cell-type r = 0.9969). **C)** Benchmark comparison with nine spatial mapping methods. Global scatter plots, analogous to panel B, summarize proportion recovery across all spot-by-cell-type pairs. Among competing methods, RCTD showed the strongest performance (r = 0.9942, ρ = 0.9071), followed by Stereoscope (r = 0.9762, ρ = 0.7544) and cell2location (r = 0.9410, ρ = 0.6508). Lower concordance was observed for DestVI (r = 0.7766), Seurat v5 (r = 0.6969), Starfysh (r = 0.6310), CellTrek (r = 0.4160), Tangram (r = 0.1137), and SPOTlight (r = -0.0406). **D)** Ground-truth dominant cell type at each spot, defined as the cell type with the highest simulated mixing proportion. The map shows a structured spatial arrangement composed of both sharply localized neuronal territories and broader supporting populations. **E)** Dominant-type maps inferred by ARCADE and competing methods. ARCADE closely recapitulates the simulated tissue organization, whereas alternative methods show varying degrees of domain simplification, boundary distortion, or local misassignment. Additional full-reference comparisons for all 22 cell types are provided in Figure S16-S18. **F)** Ground-truth spatial abundance pattern for 334 Microglia NN. The simulated proportion map spans an approximate range of 0 to 0.7, including focal enrichments and diffuse lower-level occupancy in adjacent regions. **G)** Example abundance map from cell2location, illustrating that probabilistic deconvolution can recover a broadly similar microglial territory and dynamic range but does not provide an explicit model of continuous intratype state. **H-I)** Results from discrete reference-mapping approaches, Tangram (H) and CellTrek (I). Left panels show inferred proportions, with Tangram compressing the abundance range to approximately 0–0.1 and CellTrek producing a near-binary range of 0 or 1. Right panels show the corresponding low-dimensional state-like outputs (*γ*□), which remain tied to the discrete reference representation and do not flexibly capture continuous in situ variation. **J-K)** Results from joint abundance-state generative models, Starfysh (J) and DestVI (K). Left panels show inferred abundance maps with compressed microglial ranges (0-0.3 and 0-0.23, respectively). Right panels show the corresponding latent state dimension (*γ*□ ), which varies in a way that is not cleanly aligned with the underlying spatial domain, consistent with abundance-state entanglement during joint inference. **L)** ARCADE resolves abundance-state entanglement through decoupled inference. Left, ARCADE recovers the microglial abundance landscape over the correct approximate 0-0.7 range. Right, using the inferred composition as an anchor, the spatial state model identifies a continuous latent state pattern that varies within the recovered compositional territory rather than replacing it.

We first assessed deconvolution accuracy across all 107,800 spot-by-cell-type pairs. ARCADE recovered spot-level proportions with high concordance to the simulated ground truth (Pearson r = 0.9978, Spearman ρ = 0.9193, R^2^ = 0.9954, RMSE = 0.0067, MAE = 0.0040; Figure 3B). Performance remained consistent across the reference panel, with 20 of 22 cell types showing per-cell-type Pearson r > 0.99 and a median correlation of 0.9969. We then compared ARCADE with nine widely used spatial mapping methods (Figure 3C). Among competing approaches, RCTD showed the strongest deconvolution performance (r = 0.9942, ρ = 0.9071), followed by Stereoscope (r = 0.9762, ρ = 0.7544) and cell2location (r = 0.9410, ρ = 0.6508). The comparatively strong performance of RCTD and cell2location is consistent with previous benchmarking studies identifying these methods among the more accurate tools for cell-type proportion estimation in spatial transcriptomics datasets.^29^ Methods based on discrete mapping or joint latent inference showed larger deviations from ground truth under continuous mixing conditions, including DestVI (r = 0.7766), Seurat v5 (r = 0.6969), Starfysh (r = 0.6310), CellTrek (r = 0.4160), Tangram (r = 0.1137), and SPOTlight (r = -0.0406). These differences were also evident in reconstructed spatial maps, where ARCADE more faithfully recovered both sharply localized neuronal domains and broader glial distributions across the 22-cell-type reference (Figure 3D, E; Figure S16-S18).

Because the central aim of ARCADE is not only abundance estimation but also separation of abundance from continuous cell state, we next examined a representative population with spatially graded behavior, 334 Microglia NN. In the ground truth, this population occupies a continuous abundance landscape spanning approximately 0 to 0.7, with focal enrichments and diffuse lower-level infiltration into neighboring regions (Figure 3F). This example revealed distinct classes of methodological limitation. Probabilistic deconvolution models such as cell2location, RCTD, and Stereoscope recovered abundance patterns with reasonable dynamic range, but they do not explicitly model continuous intratype state. By contrast, discrete reference-mapping tools such as Tangram and CellTrek distorted abundance scaling: Tangram compressed inferred proportions to approximately 0-0.1, whereas CellTrek produced a near-binary range of 0 or 1 (Figure 3G-I). Although their accompanying state-like outputs remained spatially coupled to these abundance estimates, those outputs remained constrained by the discrete reference representation and therefore could not flexibly describe continuous in situ variation.

We then evaluated joint generative models designed to infer abundance and state simultaneously. In both Starfysh and DestVI, the inferred microglial abundance range was compressed relative to the ground truth (0-0.3 and 0-0.23, respectively), and the corresponding latent state dimension varied in a manner that did not align cleanly with the underlying spatial territory (Figure 3J, K). These patterns are consistent with the non-identifiability problem highlighted in our controlled analyses: when abundance and state are inferred jointly from the same mixed signal, local expression differences can be absorbed by both variables. ARCADE addresses this issue by first estimating composition and then using that estimate as an abundance anchor during state inference. Under this decoupled scheme, ARCADE recovered the microglial abundance landscape over the correct 0-0.7 range while also identifying a spatially continuous latent state pattern that varied within, rather than in place of, the inferred compositional domain (Figure 3L).

To examine whether this behavior generalized beyond a single tissue and reference structure, we further constructed cross-species benchmarks using public human middle dorsolateral prefrontal cortex snRNA-seq data (donor Br2720)^30^ processed with the Seurat v5-based ARCADE pipeline. Using a 12-lineage reference, we simulated both 4,900-spot and 10,000-spot tissues Figure S19 and S20). Across both settings, ARCADE maintained high deconvolution concordance (r > 0.999, *ρ* > 0.997) and recovered spatial domains across the human reference panel (Figure S21). Competing approaches showed larger scale-dependent distortions, and in these human simulations SPOTlight and Tangram frequently converged to highly simplified spatial solutions dominated by a single lineage (Figure S19b and 20b; Figure S22-S26). Together, these benchmarks indicate that ARCADE performs favorably in structured semi-synthetic settings and, more importantly, that the decoupled design better preserves the distinction between where a cell type is enriched and how that cell type varies transcriptionally across space.

### Spatial analysis of matched human DLPFC reveals reference-stable but method-dependent tissue organization

A practical challenge in spatial transcriptomics is that the single-cell reference and spatial dataset are often derived from different donors, introducing biological and technical differences that can affect downstream deconvolution. To evaluate ARCADE in a human tissue setting while minimizing this source of variation, we used a matched multi-omic dataset generated from a single dorsolateral prefrontal cortex (DLPFC) tissue block from donor Br6432^30^. Serial sectioning of the same specimen produced closely aligned snRNA-seq, 10x Genomics Visium, and Visium Spatial Proteogenomics data (Figure 4A), allowing the single-nucleus reference to reflect the local cellular landscape of the corresponding spatial sections more directly than a cross-donor reference.

**Figure 4.**
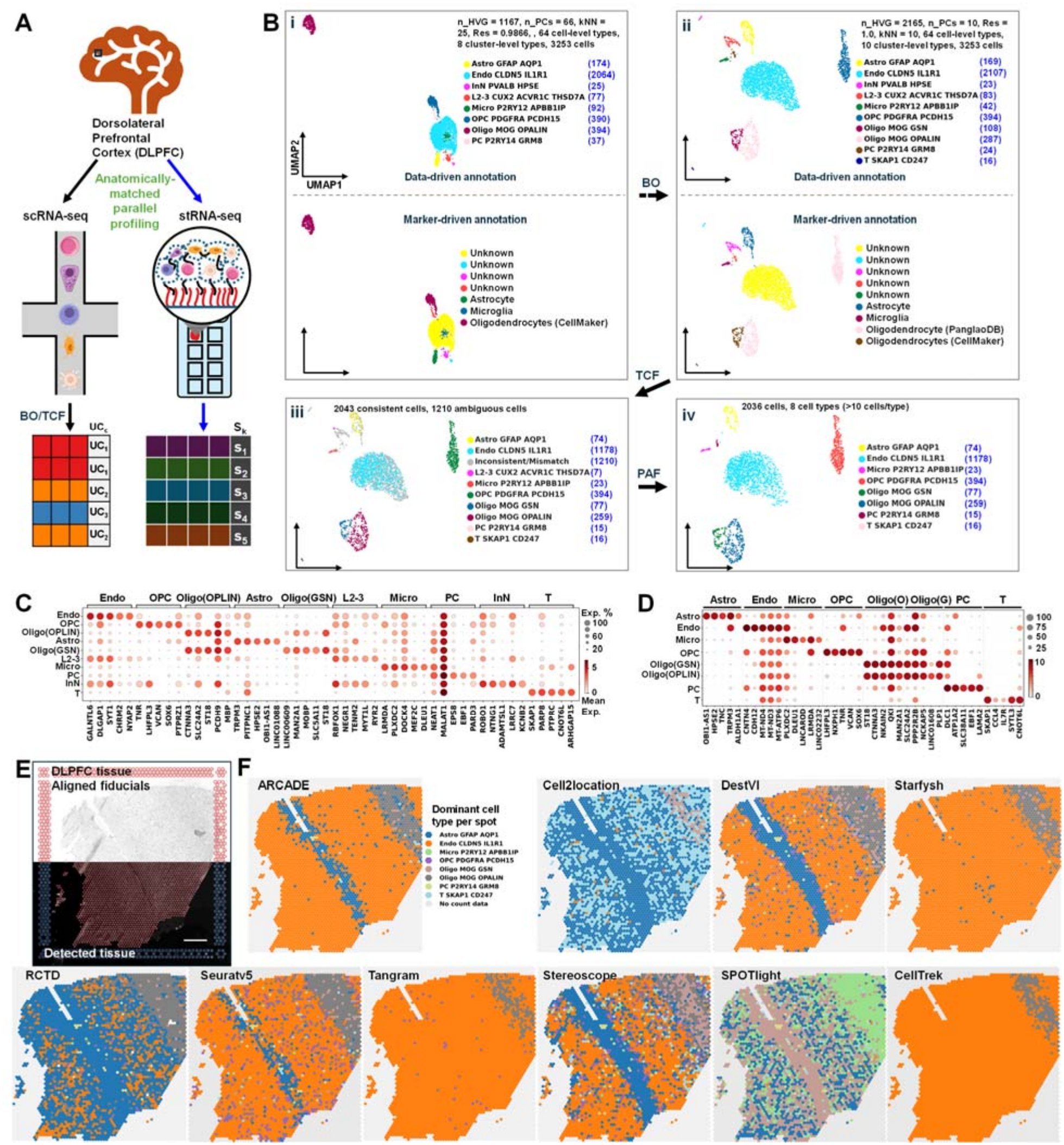
Spatially resolved analysis of the human DLPFC using CellTypist-guided reference optimization and matched-reference deconvolution. **A**) Overview of the ARCADE workflow applied to anatomically matched human anterior DLPFC tissue (donor Br6432_ant). Serial sections from the same tissue block were profiled by snRNA-seq, 10x Genomics Visium, and Visium Spatial Proteogenomics, enabling construction of a locally matched single-nucleus reference for spatial analysis. Raw snRNA-seq data were first processed by Gene Universe Harmonization (GUH) to restrict the feature space to genes shared with the paired Visium dataset. The harmonized reference then underwent Bayesian optimization over preprocessing and graph-construction parameters, followed by Tri-Consensus Filtering (TCF) and Population Abundance Filtering (PAF) to produce the final purified reference used for downstream spatial deconvolution. **B**) Bayesian hyperparameter tuning and reference purification trajectory. (i) An intermediate, sub-optimal BO trial (n_HVG = 1,167; n_PCs = 66; k-NN = 25; resolution = 0.9866) partitioning the manifold (n = 3,253 cells) into 8 broad consensus clusters. Top: Data-driven UMAP projection. Bottom: Corresponding marker-gene-driven annotation UMAP. Using an F1-score threshold of 0.1, the marker-based approach maps four distinct data-driven populations (Astrocyte [n = 174], Microglia [n = 92], OPC [n = 390], and Oligodendrocyte [n = 394]) to just three canonical cell types: astrocytes, microglia, and oligodendrocytes. It yields “Low Confidence” (Unknown) classifications for all remaining clusters. (ii) The BO-converged optimal parameter set (n_HVG = 2,165; n_PCs = 10; k-NN = 10; resolution = 1.0). Top: Optimal data-driven UMAP projection resolving 10 high-fidelity consensus clusters (revealing newly distinguished sub-populations: Oligo MOG GSN, n = 108; and T SKAP1 CD247, n = 16). Bottom: Corresponding canonical annotation UMAP exposing the severe limitations of static marker databases. Only five clusters (Astro GFAP AQP1, Micro P2RY12 APBB1IP, OPC PDGFRA PCDH15, Oligo MOG GSN, and Oligo MOG OPALIN) successfully surpass the F1 > 0.1 threshold. However, they are redundantly mapped to just four database-derived identities: astrocytes, microglia, CellMarker^31^-defined oligodendrocytes, and PanglaoDB^32^-defined oligodendrocytes. Due to failing the F1 > 0.1 confidence threshold, the remaining 5 out of 10 distinct clusters are misclassified as “Unknown.” (iii) Topological Coherence Filtering (TCF) applied to the optimal manifold removes 1,210 topologically ambiguous cells (grey), retaining exactly 2,043 highly consistent cells (colored) across 9 distinct clusters. (iv) Final purified reference achieved after Population Abundance Filtering (PAF) selectively removes extreme minority populations (n < 10). The single data-driven UMAP displays the final retained 8 robust, high-confidence cell populations (n = 2,036 cells), establishing a purified reference unaffected by the systemic biases of static marker-gene annotations. **C-D**) Cluster-level marker structure before and after ARCADE filtering. Dot plots show the top five marker genes per cluster, with color indicating normalized expression level and dot size indicating detection fraction. (C) Pre-filtered reference corresponding to the 10-cluster optimal BO solution shown in B(ii). (D) Post-filtered reference after combined TCF and PAF, showing more restricted and lineage-consistent marker expression across the retained 8-state atlas. **E**) Histological image processing and spatial registration of the paired Br6432_ant Visium section. The split-view composite shows the aligned Visium coordinate grid and fiducial landmarks (top) together with the tissue detection mask defining the analyzed region of interest (bottom). **F**) Comparative spatial reconstruction using the matched purified reference. Dominant cell-type maps are shown for ARCADE and nine comparison methods (Cell2location, DestVI, Starfysh, RCTD, Seurat v5, Stereoscope, Tangram, SPOTlight, and CellTrek). Each spot is colored according to the cell type with the highest inferred proportion among the 8 retained reference states. Although all methods used the same purified matched reference, they produced differing spatial organizations, indicating that model-specific assumptions remain an important source of variation in tissue-level interpretation.

To implement ARCADE within a Python-based analysis workflow, we used CellTypist as the annotation engine for reference optimization. As in the preceding analyses, raw snRNA-seq profiles were first subjected to GUH, restricting the reference feature space to genes also detected in the paired Visium dataset. This step reduces cross-modal feature mismatch and focuses optimization on genes measurable in the spatial assay. We then applied Bayesian optimization across the main preprocessing parameters governing reference construction, including the number of highly variable genes (HVGs), principal components (PCs), k-nearest neighbors, and clustering resolution. To first characterize intrinsic topological stability without introducing marker-based priors, we optimized a balanced objective based on CAS and MCS alone. Across a 150-trial search, broad manifold organization remained sensitive to parameter choice even though fine-grained initial labels were comparatively stable, with 64 individual cell labels observed across trials but only 3 to 11 broad consensus clusters resolved at the macroscopic level (Figure S27a; Figure S28). For biologically plausible configurations, the balanced objective ranged from approximately 63 to 73, whereas more extreme parameter combinations reduced either annotation agreement or topological consistency and were excluded from further consideration. Inspection of representative trials illustrated several recurrent failure modes. Intermediate parameterizations often produced under-resolved topologies, including manifolds with 7 or 8 broad consensus clusters (Figure S27b, c; Figure 4B, i). In the 8-cluster example shown in Figure 4B, i, marker-based annotation could confidently identify only a subset of populations, and multiple transcriptionally distinct clusters were redundantly mapped to the same broad canonical labels, while the remainder failed to pass the annotation confidence threshold and were labeled Low Confidence. These results suggest that parameter settings yielding superficially coherent embeddings may still compress biologically distinct populations into overly coarse topological states.

The data-driven search converged on an optimal configuration with a balanced score of 73.01 (2,165 HVGs, 10 PCs, k-NN = 10, resolution = 1.0), resolving 10 consensus clusters (Figure 4B, ii). We compared this optimum with other high-performing trials in the same region of parameter space, including a closely related solution differing primarily in feature selection (2,892 HVGs) and a more structurally distinct configuration (4,849 HVGs, 75 PCs) (Figure S27d, e). These comparisons indicated that nearby optima could produce similar global scores while differing modestly in how specific populations were separated or how strongly they matched static marker references. In the selected 2,165-HVG solution, only 5 of 10 mathematically coherent clusters exceeded the strict marker-based F1 > 0.1 confidence threshold (Table S1), underscoring the limited ability of static marker resources to fully validate a topology resolved from matched human tissue data alone.

Because the selected optimum remained only partially supported by marker-based validation, we next applied ARCADE’s post-optimization filtering steps to remove discordant assignments and refine the manifold into a high-confidence reference for spatial mapping. Tri-Consensus Filtering (TCF) removed 1,210 cells with discordant local, neighborhood, and globally refined identity assignments, leaving 2,043 concordant cells across 9 clusters (Figure 4B, iii). Population Abundance Filtering (PAF) then removed extreme minority populations with n < 10, eliminating the small L2-3 excitatory neuron group and yielding a final purified reference of 2,036 cells across 8 robust cell states (Figure 4B, iv). To examine whether biological priors would materially alter this endpoint, we performed a parallel optimization in which the marker prior score (MPS) was incorporated into the Bayesian objective (Figure S29; Table S2). This search converged to a distinct intermediate manifold (3,253 cells; 4,849 HVGs; 10 PCs; k-NN = 10; resolution = 0.92) and more readily flagged biologically discordant assignments as low-confidence states. However, after applying the same TCF and PAF procedure, the MPS-guided and purely data-driven routes converged on broadly similar filtered cellular cores with closely related population composition. Direct comparison of the two final filtered references showed that this convergence was nearly exact at the single-cell level: of 2,004 cells shared between the two outputs, all retained identical final annotations, with no label changes detected after consensus filtering (Figure S30). The remaining differences were limited to a small number of cells uniquely retained by one route or the other (22 unique to the MPS-guided output; 39 unique to the MPS-free output), resulting only in minor abundance shifts across the eight final populations, including Endo CLDN5 IL1R1 (1,176 versus 1,178 cells), OPC PDGFRA PCDH15 (393 versus 394), Oligo MOG OPALIN (256 versus 259), Astro GFAP AQP1 (72 versus 74), and Oligo MOG GSN (68 versus 77) ( Figure S1). Notably, inclusion of MPS did not eliminate the prominence of the cluster provisionally labeled Endo CLDN5 IL1R1, and its top marker-based matches remained weak, including neuronal or precursor-like categories with uniformly low F1 scores and minimal gene overlap (Table S3). A similar pattern was observed in the MPS-free reference, where Endo CLDN5 IL1R1 also showed only weak correspondence to external marker-defined categories, again dominated by low-scoring neuronal or precursor-like matches supported by one or two overlapping genes (Table S4). These comparisons suggest that, in this matched human cortical dataset, inclusion of biological priors influenced the intermediate optimization trajectory but had limited effect on the final filtered reference. Given the strong concordance between the MPS-guided and MPS-free purified outputs, we used the reference derived from the primary data-driven optimization for downstream spatial analyses.

To assess how filtering affected the molecular specificity of the reference, we compared cluster-level marker structure before and after TCF/PAF. Differential expression analysis showed that the pre-filtered clusters identified after Bayesian optimization were already separable at baseline (Figure 4C), but several retained broader off-target expression across neighboring populations. After filtering, marker profiles became more restricted, with stronger diagonal specificity in the cluster-by-marker dot plot and reduced low-level background across non-target groups (Figure 4D). We further compared each filtered cluster with marker gene-based matches from an external annotation resource under both the MPS-free, data-driven and MPS-guided optimization routes (Table S3 and S4). For most populations, the top-ranked database labels were concordant with the data-driven assignments, including astrocytic, T cell, pericyte, microglial, and oligodendrocyte clusters. In particular, both Oligo MOG GSN and Oligo MOG OPALIN mapped primarily to oligodendrocyte-related categories, consistent with closely related transcriptional states within the same lineage rather than distinct major cell classes. For example, in the MPS-free route, Oligo MOG OPALIN matched oligodendrocyte-related labels such as “oligodendrocytes,” “oligodendrocyte,” and “mature oligodendrocyte,” supported by recurrent overlap of canonical myelin-associated genes including OPALIN and PLP1, whereas Oligo MOG GSN also mapped chiefly to oligodendrocyte-related entries, albeit with lower scores, consistent with a related but less canonically captured oligodendrocyte state. Comparable oligodendrocyte-oriented matches were retained in the MPS-guided route, further indicating that these two clusters represent transcriptionally distinct oligodendroglia states rather than unstable reassignment across major lineages. The principal exception was the cluster provisionally labeled Endo CLDN5 IL1R1. Across both optimization schemes, its top-ranked alternative matches included neuronal or precursor-like categories, but these assignments were uniformly weak, with very low F1 scores and only one or two overlapping genes per match. We therefore do not interpret these database matches as sufficient evidence for a neuronal reannotation. Instead, they indicate that this cluster is not cleanly captured by broad static marker resources. Accordingly, we treat Endo CLDN5 IL1R1 as a provisional label and interpret this population cautiously: it retains endothelial-associated features in the data-driven manifold, but its unusually high abundance and weak external marker concordance suggest possible residual admixture, local tissue-specific expression, or incomplete representation in current marker databases. Importantly, the persistence of this same broadly defined population across both optimization settings, together with the near-complete annotation agreement between the two final filtered outputs, indicates that its recovery is not simply an artifact of whether MPS is included in the Bayesian objective. Rather, it reflects a stable reference-defined population whose precise canonical identity remains unresolved under current marker-based frameworks. Together, these analyses indicate that ARCADE’s filtering steps improve practical specificity for most populations while also identifying clusters whose lineage assignment remains provisional.

We next applied the resulting reference to the paired Visium section after image processing and coordinate registration (Figure 4E). Using the same purified 8-state reference for all methods, we compared ARCADE with nine widely used spatial deconvolution approaches by assigning each spot to its dominant inferred cell type (Figure 4F). Despite operating on the same input reference, the methods produced notably different tissue-level architectures. ARCADE resolved three broad spatial domains: a diagonal region enriched for astrocytes, a right-corner domain enriched for oligodendrocytes, and a broader background territory with substantial signal from the provisionally labeled Endo CLDN5 IL1R1 population. Similar large-scale structure was partially recapitulated by DestVI, Seurat v5, and Stereoscope, whereas agreement across methods was less consistent for individual fine-scale territories. The right-corner OPALIN-positive oligodendrocyte domain was detected by most methods except SPOTlight, while the diagonal astrocyte stripe and extended CLDN5 IL1R1-associated territories showed more method-dependent recovery. These differences indicate that, even under a shared matched-reference setting, the computational assumptions of the deconvolution model continue to shape the inferred spatial organization of human cortical tissue.

### ARCADE explicitly resolves spatial intratypic heterogeneity and decouples cellular abundance from state

Having established a matched reference for the DLPFC sections, we next used the paired Visium and Visium-SPG data to test whether ARCADE could separate physical cell enrichment from localized transcriptional activation. The aligned protein measurements provided an orthogonal spatial benchmark for evaluating abundance estimates in situ. We first examined sample Br6432 using the GFAP immunofluorescence (IF) channel as an external spatial reference for astrocyte-associated signal. The isolated GFAP channel revealed two prominent domains of elevated signal: a broad diagonal band and a second localized region in the right corner of the tissue (Figure 5A). The corresponding spatial transcript map of GFAP mRNA recapitulated this pattern (Figure 5B), indicating that both protein and transcript measurements captured the same broad anatomical organization. We therefore asked whether deconvolution methods would interpret these two regions similarly in terms of astrocyte abundance.

**Figure 5.**
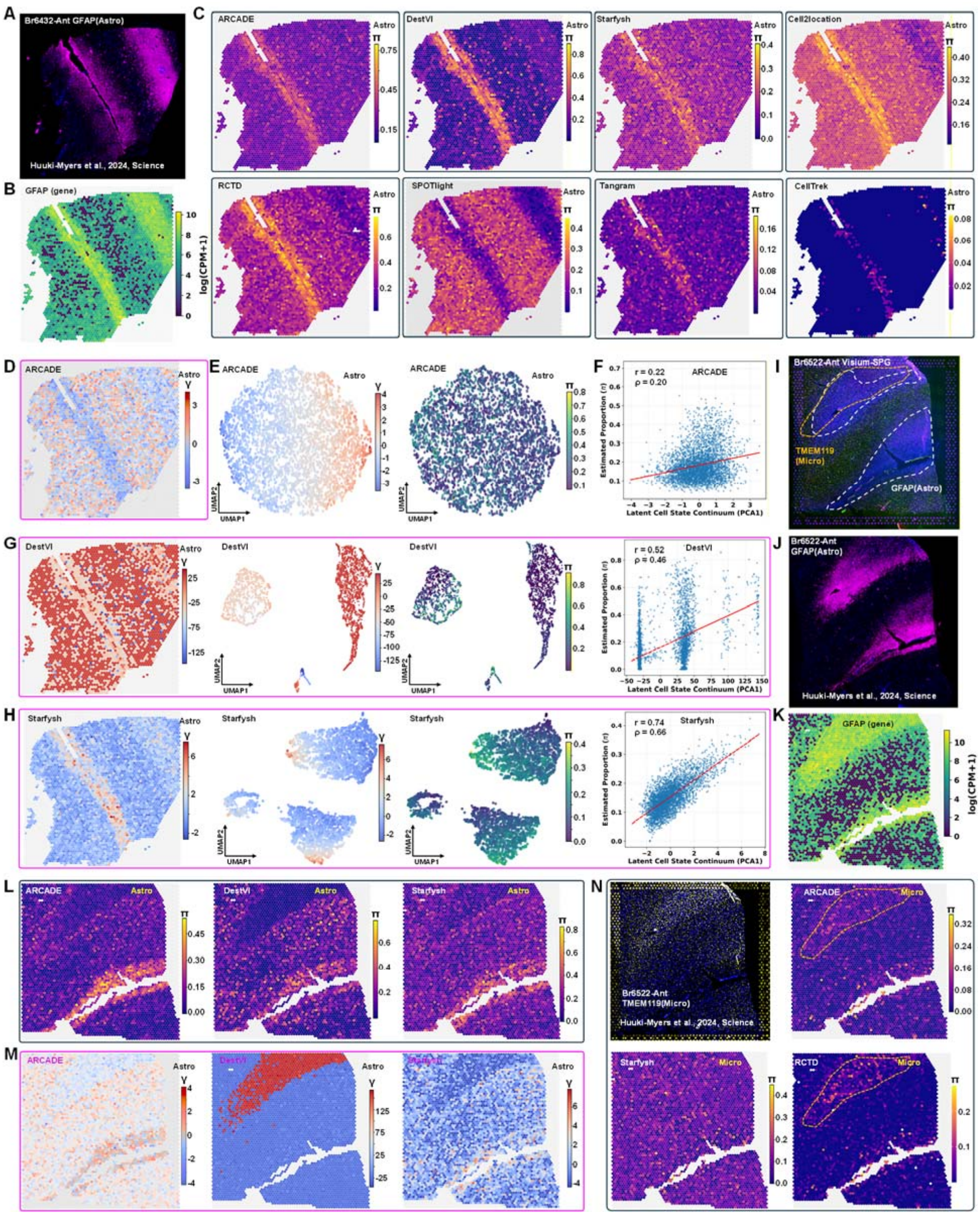
Orthogonal proteogenomic validation of abundance-state decoupling in matched human DLPFC sections. **A)** Single-channel GFAP immunofluorescence (IF) image from Visium Spatial Proteogenomics (Visium-SPG) for a matched human DLPFC section (sample Br6432). The isolated GFAP channel shows two astrocyte-associated domains: a broad diagonal band and a localized right-corner region. The image was rendered from publicly available raw Visium-SPG data from Huuki-Myers et al. (2024); the corresponding multiplexed visualization is shown in their Figure S28. **B)** Spatial feature plot of GFAP mRNA expression (log(CPM+1)) across the paired Br6432 Visium section. **C)** Spatial maps of inferred astrocyte proportions in Br6432 for ARCADE and baseline deconvolution methods. Color scale denotes estimated astrocyte fraction per spot. **D)** Spatial map of astrocyte transcriptional states inferred by ARCADE in Br6432, highlighting three broad regimes corresponding to the diagonal band, right-corner domain, and background tissue. **E)** Left, UMAP of ARCADE-inferred astrocyte latent states in Br6432, where each point represents a Visium spot positioned according to its deconvolved astrocyte-specific transcriptomic state. Right, the same embedding colored by inferred astrocyte abundance, allowing visual comparison between state variation and local proportion. **F)** Relationship between astrocyte state and abundance in Br6432. The x axis shows the first principal component of the astrocyte latent variable *γ*, and the y axis shows the inferred astrocyte proportion *π*. Pearson and Spearman statistics quantify state-abundance coupling. **G-H)** Comparison with alternative state-aware models in Br6432. For DestVI (G) and Starfysh (H), panels show: spatial map of inferred astrocyte states, UMAP of the corresponding latent representation, the same UMAP colored by inferred astrocyte abundance, and the association between the dominant latent axis and astrocyte proportion. **I)** Multiplexed Visium-SPG IF image from a second matched DLPFC section (sample Br6522), showing GFAP-positive astrocyte-associated bands and a TMEM119-positive microglia-enriched band. Image rendered from publicly available raw data from Huuki-Myers et al. (2024); see their Figure S28 for the corresponding multiplexed view. **J)** Isolated GFAP IF channel from the Br6522 section, showing two parallel astrocyte-associated bands. **K)** Spatial feature plot of GFAP mRNA expression (log(CPM+1)) across the paired Br6522 Visium section. **L)** Spatial maps of inferred astrocyte proportions in Br6522 for ARCADE, DestVI, and Starfysh. **M)** Spatial maps of inferred astrocyte transcriptional states in Br6522 for ARCADE, Starfysh, and DestVI. **N)** Top left, isolated TMEM119 IF channel from Br6522, marking a microglia-enriched anatomical band. Remaining panels show inferred microglia proportions for the same section from ARCADE, Starfysh, and RCTD.

Across methods, the dominant diagonal astrocyte-associated region was recovered relatively consistently, including by ARCADE, cell2location, RCTD, and Seurat v5 (Figure 5C; Figure S31). In contrast, the right-corner domain showed substantially greater method-dependent variability. Several approaches inferred comparatively low astrocyte abundance in this region despite the strong GFAP protein and mRNA signal, whereas CellTrek recovered the general two-domain pattern but with low overall astrocyte proportions. These differences are consistent with a setting in which elevated GFAP expression does not necessarily reflect a proportional increase in local astrocyte number.^33, 34^ In the DLPFC, increased GFAP signal can accompany reactive or state-shifted astrocytes, suggesting that methods that couple abundance estimation too tightly to marker intensity may misinterpret local activation as density, or conversely undercall abundance when the marker program departs from the reference average. ARCADE was designed to address this ambiguity by first anchoring cell-type composition and then modeling continuous intratype variation conditional on that compositional estimate. In Br6432, this yielded a spatial astrocyte state map with three broad regimes: the diagonal band, the right-corner domain, and the remaining background tissue (Figure 5D). Importantly, the right-corner region was represented primarily as a distinct astrocyte transcriptional state rather than simply as a second high-abundance peak. This interpretation is consistent with the view that part of the GFAP elevation in this region reflects altered per-cell expression rather than only increased astrocyte density.

To visualize this separation more directly, we projected the inferred astrocyte latent representations onto a UMAP manifold (Figure 5E, left) and overlaid the corresponding abundance estimates on the same embedding (Figure 5E, right). The latent space formed a broad continuum rather than a set of sharply isolated groups, consistent with graded intratype variation. Abundance varied across this continuum, but not in a way that collapsed the manifold into a simple density axis. Quantitatively, the dominant axis of astrocyte state variation showed only a weak association with inferred abundance (Pearson r = 0.22; Spearman ρ = 0.20; Figure 5F), supporting the interpretation that ARCADE can capture state variation without forcing a one-to-one mapping between activation and local proportion. Extending the same analysis across all eight reference cell types yielded similarly weak state-abundance coupling with substantial residual dispersion, suggesting that this behavior is not restricted to astrocytes (Figure S32).

We next compared ARCADE with methods that also model continuous or semi-continuous cellular variation. DestVI identified the right-corner region as distinct from background tissue, broadly agreeing that this domain is transcriptionally unusual (Figure 5G). However, its latent embedding was more fragmented and exhibited stronger state-abundance coupling (ρ = 0.46), suggesting greater leakage between the inferred composition and state dimensions. Starfysh likewise separated major spatial domains (Figure 5H), but its inferred astrocyte states showed substantially stronger coupling to abundance (ρ = 0.83) and lower residual dispersion, consistent with a latent representation that tracked density more directly. Across all eight cell types, these trends persisted: ARCADE generally produced broader continuous manifolds with weaker state-abundance coupling, whereas DestVI and Starfysh more often yielded discretized or abundance-associated latent structures (Figure S33 and S34). We also examined Tangram and CellTrek for spatial state mapping (Figure S35). In this setting, Tangram left a subset of spots effectively unresolved, whereas CellTrek showed limited within-type state structure, indicating that these approaches were less suitable for representing continuous intratype variation in this sample.

Because ARCADE depends on the upstream reference, we next asked whether these observations were robust to the reference-generation ecosystem. In addition to the CellTypist-based reference used above, we constructed an independent reference using a Seurat-based pipeline and optimized its preprocessing parameters by Bayesian search. As in the CellTypist setting, heuristic parameter choices produced marked changes in topology and annotation coherence, whereas the optimized solution converged to a stable four-lineage reference comprising Pax6, oligodendrocyte, OPC, and astrocyte populations (Figure S36 and S37). Applying TCF removed 343 topologically ambiguous cells and retained 2,910 concordant cells for downstream mapping. The filtered reference showed sharper cluster-restricted marker profiles and reduced cross-cluster contamination, providing an independent test bed for spatial analysis. Notably, one major reference population remained difficult to interpret unambiguously. Using this Seurat-derived reference, we repeated the Br6432 analysis. The matched GFAP IF and transcript maps again showed the same two broad astrocyte-associated domains (Figure S38a and 38b). ARCADE assigned high astrocyte abundance to the primary diagonal region while maintaining comparatively modest abundance in the right-corner domain (Figure S38c), again suggesting that part of the right-corner signal is better explained by transcriptional state than by cell number alone. In latent space, ARCADE recovered broad continuous manifolds with weak state-abundance coupling across all four lineages (Figure S39); for astrocytes, the association was close to zero (Pearson r = 0.012; Spearman ρ = 0.068). DestVI and Starfysh showed the same qualitative limitations observed with the CellTypist reference, including fragmented latent structure and stronger abundance dependence (Figure S40 and S41). Together, these results indicate that ARCADE’s decoupling behavior is reproducible across two independently optimized reference construction pipelines.

We then asked whether the same conclusions held in an independent tissue section, Br6522. In this sample, Visium-SPG imaging revealed a more complex architecture: the GFAP channel showed two parallel astrocyte-associated bands, while the TMEM119 channel marked a distinct microglia-enriched band in the upper region of the section (Figure 5I and 5J). The spatial transcript map of GFAP closely matched the dual-band protein pattern (Figure 5K). When we compared inferred astrocyte abundance across methods, several approaches assigned little or no astrocyte signal to one or both validated bands, whereas ARCADE, cell2location, and Starfysh more clearly recovered the upper domain (Figure 5L; Figure S42). ARCADE further resolved three broad astrocyte state regimes corresponding to the upper band, lower band, and background tissue (Figure 5M), indicating that the model could represent multiple spatially organized astrocyte programs within the same section. As in Br6432, latent-space analysis in Br6522 supported weak coupling between state and abundance across major neural lineages (Figure S43). By comparison, DestVI again tended to produce more fragmented embeddings (Figure S44), while Starfysh showed stronger linear dependence between inferred states and abundance with lower residual dispersion (Figure S45). These sample-to-sample comparisons suggest that ARCADE’s two-stage structure is comparatively stable across tissues with different spatial topologies, whereas joint inference can be more sensitive to how marker activation and composition co-vary across the section.

Finally, we examined whether orthogonal protein measurements also supported ARCADE’s abundance estimates for another lineage. In Br6522, the isolated TMEM119 IF channel marked a clear microglia-enriched anatomical band (Figure 5N, top left). ARCADE and RCTD both recovered elevated microglial abundance in this same region, whereas several other methods returned flatter spatial patterns or markedly attenuated signal (Figure 5N; Figure S46). This complementary result suggests that the matched proteogenomic data are informative not only for astrocyte-associated state-abundance separation, but also for evaluating abundance recovery in lineages with more localized spatial enrichment. Together, these analyses support the view that ARCADE can separate spatial abundance from continuous intratype transcriptional variation in human cortex while remaining robust to alternative reference construction backbones and distinct anatomical contexts.

## DISCUSSION

Spatial transcriptomics is increasingly used to reconstruct tissue organization from measurements that conflate multiple biological sources of variation within each spot. A central difficulty is that changes in spot-level gene expression can reflect either differences in cell-type abundance or continuous changes in transcriptional state within a shared cell type. When these two factors are inferred jointly without sufficient constraint, localized state variation can be absorbed into abundance estimates, complicating biological interpretation. ARCADE was developed to address this problem by anchoring spatial inference to a fixed compositional prior and then modeling residual, within-cell-type transcriptional variation as a separate latent component. Across controlled benchmarking and matched human tissue analyses, this two-stage formulation improved the separation of composition from state and yielded spatial patterns that were more consistent with orthogonal histological and molecular features.

An additional challenge in many spatial workflows is the dependence on single-cell references assembled across donors, protocols, or studies. Large integrated atlases are valuable for defining broad cell taxonomies, but they can also introduce biological and technical mismatches between reference and target tissue. Such mismatches are especially relevant when the goal is not only to recover major lineages, but also to detect localized and potentially subtle transcriptional variation within them. In this setting, an anatomically matched reference can provide a practical advantage by reducing cross-sample discordance before deconvolution is performed. Our analyses in human DLPFC support this view: using matched single-cell references allowed us to evaluate spatial abundance and state inference without the additional uncertainty introduced by donor-to-donor integration. At the same time, our results also show that reference matching alone is not sufficient. Even with an anatomically aligned reference, inferred spatial organization remained sensitive to how the reference was constructed and how abundance and state were modeled.

These observations motivated the upstream reference-optimization component of ARCADE. In practice, standard single-cell annotation pipelines can be highly sensitive to heuristic choices such as feature selection, clustering resolution, and neighborhood graph construction. We therefore used Bayesian optimization and consensus-based filtering to identify reference configurations with improved topological stability and annotation coherence before spatial mapping. Importantly, we do not view this step as the primary conceptual advance of the method. Rather, it serves as an enabling layer that reduces avoidable instability in the reference and makes downstream abundance-state decoupling more reliable. The matched DLPFC analyses also illustrate an important limitation: even after optimization, not every recovered population can be assigned a definitive canonical identity. In particular, some reference-defined clusters showed weak or conflicting marker-based matches, indicating that improved topological consistency does not automatically resolve biological annotation ambiguity. For this reason, we interpret such populations conservatively and emphasize that reference optimization should be understood as improving robustness, rather than guaranteeing perfect cell-type labeling.

A useful implication of this framework is that integrated and sample-specific references need not be treated as competing strategies. Integrated atlases remain well suited for establishing broad cross-donor taxonomies and for defining stable major lineages across cohorts. Sample-matched references, by contrast, may be preferable when the main objective is to preserve local biological variation and minimize distortions introduced by integration. ARCADE is compatible with both settings. In one use case, a broad integrated reference could define the compositional scaffold, after which ARCADE models localized intratype transcriptional variation within the target tissue. In another, a matched single-donor reference can be used directly when donor concordance is a priority. We anticipate that the most informative applications will often combine these modes: stable lineage definitions from integrated references, together with localized state inference that remains sensitive to tissue-specific biology.

Our comparison of CellTypist- and Seurat-based reference pipelines further suggests that the framework is not tied to a single annotation ecosystem. In our hands, the two backbones exhibited different practical behaviors on the same human DLPFC material. The CellTypist-based workflow tended to preserve finer transcriptomic subdivisions, which may be useful when the goal is to retain rare or spatially restricted populations. The Seurat-based workflow produced a more consolidated set of broad lineages, which can be advantageous when a stable coarse-grained tissue architecture is desired. We do not interpret these differences as a universal ranking of platforms; rather, they indicate that reference construction remains partly objective-dependent. A key feature of ARCADE is therefore not allegiance to one backend, but the ability to place distinct reference-generation strategies into a common downstream framework for spatial abundance-state decoupling.

Overall, ARCADE supports a view of spatial transcriptomics in which cellular composition and transcriptional plasticity should be modeled as related but distinct sources of variation. By combining reference optimization with anchored two-stage inference, the framework seeks to improve the interpretability of spatial mapping in heterogeneous tissues, particularly in settings where localized state changes might otherwise be mistaken for changes in cell abundance. We anticipate that this distinction will be especially important in contexts such as neuroinflammation, tissue remodeling, and tumor ecosystems, where microenvironmental cues can reshape canonical marker expression without corresponding changes in cell number.

## Methods

### Automated CellTypist based scRNA-seq Processing, BO, and TCF

#### Data Ingestion and Spatial Transcriptomic Alignment

The analytical pipeline was implemented in Python 3.9+ utilizing Scanpy (v1.9+), CellTypist (v1.6+), scikit-optimize (v0.9+), and harmonypy (v0.0.9+). To ensure seamless integration with diverse single-cell workflows, the pipeline natively ingested multiple data formats, including 10x Genomics directories, HDF5 matrices (.h5), and preprocessed AnnData objects (.h5ad). Crucially, to ensure strict feature compatibility for downstream spatial deconvolution, a global spatial intersection was executed immediately following data ingestion. The comprehensive scRNA-seq feature space was strictly filtered to retain only genes actively detected within the paired ST reference matrix.

A rigorous quality control (QC) protocol was applied across all datasets. Low-quality cells and suspected multiplets were excluded by filtering events expressing fewer than 200 or more than 7,000 unique genes. Cells exhibiting stress signatures were discarded based on a maximum mitochondrial read fraction of 10%, with mitochondrial transcripts identified dynamically via regular expression matching of standard prefixes (MT-, Mt-, mt-followed by hyphens, periods, underscores, or colons). Sparsely expressed features were eliminated. The surviving count matrix was subjected to library size normalization to 10,000 counts per cell (CP10k) and log (1 + *x*) transformed. This normalized profile was preserved in the object’s .raw state to serve as the definitive foundation for downstream probabilistic annotation and differential gene expression analysis.

#### Topological Pipeline and Parameter Space Encoding

To systematically explore the hyperparameter landscape governing single-cell dimensionality reduction and clustering, each candidate configuration was mathematically encoded as a four-dimensional vector: *x* = [*n*_*HVG*_,*n*_*PCs*_,*n*_*neighbors*_, resolution]. The bounded search space was defined as *n*_*HVG*_ *∈* [800,20000](dynamicayl upper-bounded by available genes), *n*_*PCs*_ *∈* [10,100], *n*_*neighbors*_ *∈* [10,50], and resolution *∈* [0.2,1.0].

Highly variable genes (HVGs) were selected using one of two modes: (1) a standard rank-based approach (Seurat v3 flavor) selecting the top *n*genes by normalized dispersion, or (2) a highly stringent two-step sequential filtration that first bounded genes by mean expression and minimum dispersion before ranking. For multi-sample cohorts, HVG selection was executed in a batch-aware manner.

The expression matrix was subset to the selected *n*_*HVG*_, Z-score scaled (capped at 10), and subjected to Principal Component Analysis (PCA). To guarantee algorithmic convergence during iterative sub-clustering, the number of computed PCs was dynamically bounded by the matrix rank (min (*n*_*PCs*_, *N*_*cells*_ *−* 1, *N*_*genes*_ *−* 1)). For multi-sample cohorts, PCA embeddings were corrected for technical batch effects utilizing the Harmony integration algorithm, which iteratively adjusts cell positions in PCA space to remove batch-specific clustering while preserving biological variation. The corrected embedding (X_pca_harmony) was utilized for all downstream k-nearest neighbor (KNN) graph construction and Leiden community detection.

#### Composite Objective Function for Manifold Evaluation

To evaluate the structural and biological fidelity of the transcriptomic manifold, we engineered a composite objective function, *f*(*x*), synthesizing biological purity, transcriptional authenticity, and topological separation.

##### 1. Cluster Annotation Score (CAS)

Baseline cellular annotations (lj for cell j) were generated by projecting the complete transcriptomic matrix through a pre-trained probabilistic CellTypist model. For any discrete cluster C, the consensus identity L_c_was defined as the majority annotation. The localized CAS was calculated as the proportion of cells structurally aligned with this consensus:

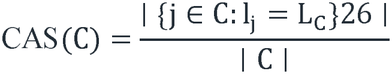

where ∣ C ∣ Represents the total cellular volume of the cluster. We subsequently derived two global metrics. The Weighted CAS (CAS_W_) represents the absolute percentage of the total cellular population (N) matching their respective topological consensus, defined mathematically as:

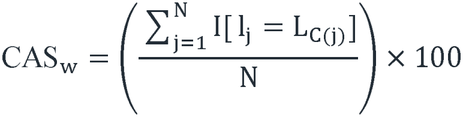

where Idenotes the indicator function yielding 1 if cell jaligns with its cluster consensus, and 0 otherwise. The Simple CAS (CAS_s_) was aggregated via two modes: leiden (the unweighted arithmetic mean across individual topological clusters, 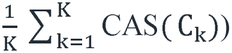 or consensus (merging disjoint clusters sharing an identical consensus label prior to averaging, thereby preventing score inflation when multiple distinct clusters represented the same overarching cell type).

##### 2. Marker Concordance and Capture Scores

Canonical markers (denoted as m_1_,…, m_n_) for each assigned cell type Twithin cluster Cwere derived via a one-vs-rest Wilcoxon rank-sum test on the unscaled full-transcriptome matrix. The Marker Concordance Score (MCS) quantified the penetrance of these markers, defined as the average proportion of cells within the cluster exhibiting positive log-normalized expression of the top ncanonical markers. Mathematically, the localized score is formulated as:

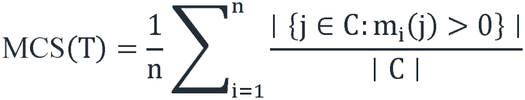

where the condition m_i_(j) > 0 indicates the positive transcriptomic detection of marker iwithin cell j. The global MCS was computed as the arithmetic mean across all qualifying cell types (strictly omitting biologically sparse populations comprising fewer than two cells to guarantee statistical validity). For manual annotation validation, a distinct Marker Capture Score was additionally computed to quantify the proportion of canonical reference markers (derived from external databases such as CellMarker) successfully recovered among the top differentially expressed genes.

##### 3. Topological Silhouette Score

To assess geometric cohesion, the silhouette score *S ∈* [−1,1] was computed on the latent space and linearly scaled to *S*_*scaled*_ *∈* [0,1], Topologies that failed to resolve heterogeneity (yielding a singular global cluster) or exhibited mathematically degenerate over-clustering (*N*_*clusters*_ *≥ N*_*cells*_) were severely penalized by forcibly setting *S*_*scaled*_ = 0 .

###### Mathematical Synthesis

Depending on the analytical paradigm, *f*(*x*)was dynamically formulated across three optimization models. The *biological* model (default) computed the geometric mean of the biological metrics; the *structural* model incorporated spatial embedding geometry; and the *silhouette* model optimized solely for cluster separation. For models integrating *k* normalized biological metrics *m*_*i*_ *∈ {CAS*_*W*_, *CAS*_*s*_, *MCS, S*_*scaled*_ *}*, the function was defined as:

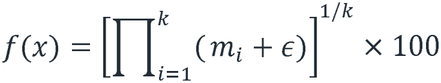

where *ε* = 10 ^−6^ ensured numerical stability,

##### 4. Marker Prior Score (MPS) via F1-based Evaluation

To rigorously benchmark the biological validity of derived cell states against established external knowledge, we introduced the Marker Prior Score (MPS), formulated as a harmonized Fl-score metric, For each inferred cluster *C*, data-driven differentially expressed genes (DEGs) were identified via a one-vs-rest Wilcoxon rank-sum test. To prioritize biologically robust markers, DEGs were ranked using a custom composite scoring system. This composite score integrated min-max normalized values of log2 fold-change (*S*_*fc*_), mean expression magnitude (*S*_*expr*_), and the difference in cellular detection percentage between the target cluster and the reference (*S*_*pct*_). The final rank score was computed as a weighted sum:

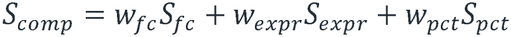

where *W*_*fC*_, *w*_*expr*_, and *w*_*pct*_ default to 0.4, 0.3, and 0.3, respectively. The top *N* (default TV = 50) highest-ranking DEGs constituted the data-driven marker set, *D*_*c*_. Concurrently, the consensus annotation for cluster *C* was mapped against a comprehensive external prior marker database. To mitigate nomenclature discrepancies, a hierarchical robust matching algorithm was deployed, sequentially evaluating: (1) biological abbreviation expansion, (2) exact and normalized string matching, (3) canonical synonym mapping, and (4) fuzzy string matching bounded by a strict Jaccard-similarity threshold (default 0.6). Successful mapping yielded a canonical marker set, *M*_*T*_, for the matched cell type *T*. Gene nomenclatures across both sets were strictly harmonized by species conventions (e.g., uppercase for human).

The localized MPS for cluster *C* was defined as the harmonic mean (F1 score) of marker precision (*P*) and recall (*R*):

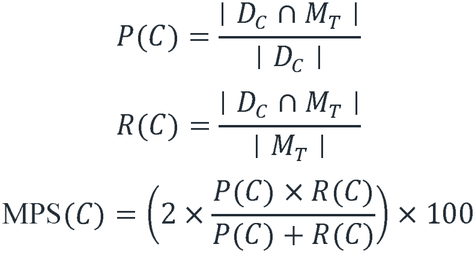

To enforce strict biological fidelity, clusters whose annotations failed to match any established reference entity were heavily penalized by assigning MPS(*C*) = 0. The global MPS was computed as the arithmetic mean of MPS(*C*) across all clusters possessing sufficient cellular volume (default ≥ 5cells).

When utilizing external biological priors, the global MPS was integrated into the Bayesian optimization framework as an additive mathematical bonus to the primary objective function *f(x)*. The final objective state was defined as:

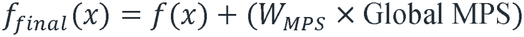

where *W*_*MPS*_ represents a predefined bonus coefficient (default 0.20), permitting up to a 20% quantitative enhancement for topologies demonstrating perfect concordance with established canonical marker profiles. Furthermore, during the initial feature space definition, the algorithm selectively shielded identified canonical markers from variance-based filtration, dynamically expanding the HVG pool to guarantee their representation in the latent embedding.

#### BO and Acquisition Strategies

Optimization was driven by Gaussian Process (GP) surrogate modeling (gp_minimize) utilizing a Matern kernel (*v* = 5/2) and Automatic Relevance Determination (ARD). The surrogate model probabilistically approximates the objective landscape as *f(x) ∼ GP(µ (x) σ* ^2^ (*x*)), where *µ (x)*is the predictive mean and σ(*x*)is the predictive standard deviation representing epistemological uncertainty at candidate configuration *x*.

To systematically navigate the multi-dimensional search space and balance the trade-off between exploration and exploitation, candidate hyperparameters were iteratively selected by maximizing three parallel acquisition strategies:

##### 1. BO-EI (Expected Improvement)

Calculates the expected magnitude of potential gains over the current global optimum score (*f*^+^), mathematically formulated as:

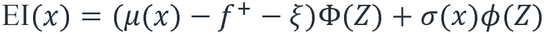

where 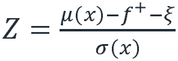, and Φ and *ϕ* represent the cumulative distribution and probability density functions of the standard normal distribution, respectively. The trade-off parameter was empirically fixed at ξ = 0.0l for balanced traversal.

##### 2. Exploit (Probability of Improvement)

Strictly prioritizes localized refinement by emphasizing the absolute statistical likelihood of surpassing the current optimum, defined as:

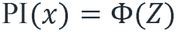

utilizing a highly exploitative trade-off parameter of ξ = 0.01.

##### 3. Explore (Exploration-weighted EI)

Utilizes the standard EI mathematical architecture but aggressively elevates the trade-off parameter to ξ = 0.10. This actively penalizes the iterative sampling of hyperparameter configurations adjacent to the current optimum, compelling the GP to investigate regions of high predictive uncertainty to escape local optima.

During each sequential iteration, the configuration yielding the highest global score across these parallel acquisition evaluations was selected for downstream pipeline execution.

#### Automated Iterative Refinement and Quality Reporting

To resolve transcriptomic ambiguities in heterogeneous sub-lineages, the pipeline executed an automated, recursive refinement protocol on subpopulations exhibiting a CAS below a predefined confidence threshold (e.g., 70%). Localized BO was re-initiated to establish microenvironment-optimal hyperparameters. Stringent dynamic stopping criteria terminated refinement if: (1) all updated clusters surpassed the CAS confidence threshold; (2) the remaining subset fell below a minimum threshold (A = 100 cells); (3) the local feature space lacked sufficient transcriptional variance (<200 detectable HVGs); or (4) degenerate over-clustering occurred, defined empirically as *N*_*clusters*_ *> N*_*ce*_*u*_*s*_*/5*.

During refinement, cumulative UMAP projections were continuously generated, contrasting cells with successfully refined annotations against those remaining below the CAS threshold (highlighted in grey). Upon completion, an automated cell-type journey summary report was synthesized, tracking each population’s total cell count, matching predictions, and aggregated CAS across all analysis stages to quantify iterative annotation improvements.

#### TCF and Deconvolution Reference Export

To ensure the absolute fidelity of the reference signature matrix supplied to spatial deconvolution algorithms, we implemented a strict TCF mechanism. This step exclusively isolated cells exhibiting unanimous annotation agreement across three distinct analytical tiers: (1) the independent per-cell probabilistic prediction; (2) the global baseline cluster consensus; and (3) the locally resolved prediction following iterative refinement.

Cells demonstrating discordance across these axes were explicitly discarded as suspected multiplets or ambiguous transitional states. This exclusion was visually validated via a Consistency Context UMAP, which plotted failing cells in grey against the validated manifold. To prevent statistical convergence errors during spatial deconvolution, a compositional filter was subsequently applied, entirely removing rare cell populations comprising fewer than a user-defined threshold of cells. Finally, the pipeline exported three perfectly dimensionally aligned CSV matrices (sc_counts.csv, sc_labels.csv, and st_counts.csv) to serve as the empirical foundation for spatial mapping.

### Automated Seurat-Based scRNA-seq Processing, BO, and TCF

#### Data Ingestion and Cross-Modality Alignment

To provide a complementary, highly versatile analytical framework, a parallel pipeline was implemented in R (v4.0+) utilizing the Seurat v5.0+ architecture and standard Bioconductor annotation databases. Data ingestion natively accommodated diverse formats, including 10x Genomics MTX, HDF5, and serialized Seurat objects, with ENSEMBL identifiers systematically mapped to standard gene symbols via org.Hs.eg.db or org.Mm.eg.db.

Quality control filtration, library size normalization (LogNormalize), and the cross-modality global spatial intersection were executed utilizing identical biological thresholds and topological logic as the Python framework (strictly enforcing bounds of 200-7,000 unique features and <10% mitochondrial transcripts).

#### Parameter Space and Feature Selection

The four-dimensional hyperparameter search space (*x*) and its boundary constraints were parameterized identically to the aforementioned Python framework. Highly variable gene (HVG) selection was executed via Seurat’s variance-stabilizing transformation (vst), dynamically toggling between the standard rank-based approach and the highly stringent, rigid numerical boundary filtration (mean and dispersion) described previously. Following Z-score scaling, Principal Component Analysis (PCA) was computed, with the number of components dynamically bounded by min (*n*_*PCs*_, *N*_*cells*_ *−* 1, *N*_*genes*_ *−* l,105)to ensure matrix stability during downstream algorithmic operations.

#### Cell Type Annotation via Reference Anchoring

Rather than utilizing pre-trained machine learning classifiers (e.g., CellTypist), baseline cellular identity assignment in the R pipeline was executed dynamically using Seurat’s Anchor-based Label Transfer algorithm (FindTransferAnchors and TransferData). During each optimization trial, the query dataset was projected into the PCA-reduced subspace of a fully annotated, user-provided reference object.

Mutual nearest neighbors (MNNs) established biological anchors between datasets. To classify query cell *q*, the algorithm computed a probabilistic prediction score *S(l)*_*q*_ for each reference cell type *l*. This was derived via an *L2*-normalized, distance-weighted transition probability matrix between the query cell and its *k*-nearest reference anchors (where *k* dynamically matched the currently evaluated n_*neighbours*_ hyperparameter). The prediction score is formally defined as:

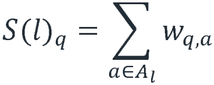

where 4_;_represents the set of anchors belonging to cell type *l*, and *w*_*q*_,_*a*_ represents the assigned transition weight between query cell qand anchor *a*. The cell type yielding the maximum prediction score was assigned as the independent individual prediction (ctpt_individual_prediction).

#### Mathematical Synthesis of the Objective Function

The optimization landscape was driven by the identical tripartite composite objective function *f(x)* established in the Python pipeline. The CAS and Topological Silhouette Score were calculated using the exact mathematical formulations described previously. For the Marker Concordance Score (MCS), candidate markers were equivalently derived utilizing Seurat’s FindAllMarkers function (Wilcoxon rank-sum test), strictly filtered for positive expression (log _2_ FC > 0). Aggregation of these metrics via the bounded geometric mean was performed identically, substituting a continuous ε = 10^−4^to ensure strict numerical stability during R-based exponentiation.

#### Fault-Tolerant BO

Hyperparameter exploration was orchestrated via the rBayesianOptimization package. While sharing the fundamental Gaussian Process (GP) architecture and acquisition strategies of the Python framework, the R implementation required distinct algorithmic safeguards against surrogate model collapse:

##### GP Covariance Stabilization

Single-cell topological evaluations frequently yield plateaued, mathematically identical fitness scores across iterations, which severely compromise standard GP optimization by producing singular, non-invertible covariance matrices. To preempt optimization failure, we engineered a deterministic stochastic jitter injection protocol. When variance among historical scores fell below 10^−10^, a mathematically negligible fractional scalar, formulated as *δ ∼ U*(10^−10^,10^−8^), was appended to the raw objective score. This explicit modification strictly preserved matrix invertibility while retaining absolute score rank order.

##### Deterministic Fallback Architecture

In instances of complete surrogate model failure, the script executed a pre-seeded programmatic fail-safe. Rather than terminating, the pipeline automatically transitioned from Bayesian to pseudo-random coordinate search, drawing from a pre-generated array of valid parameter geometries to fulfill the iteration quota (*N*_*ca*_*u*_*s*_), guaranteeing successful pipeline execution regardless of spatial complexity.

#### Iterative Refinement and TCF

The automated recursive refinement of low-confidence subpopulations (<70% CAS) followed the identical programmatic logic detailed above. Prior to exporting the spatial deconvolution reference, the manifold was subjected to dynamic Consistency Filtering to maintain mathematical congruence across datasets. The algorithm dynamically toggled between two stringency modes:

##### Tri-Consensus Mode

Activated if recursive refinement was deployed. This mode exclusively isolated cells demonstrating unanimous agreement across three vectors: *I(c) = C*_*ind*_ (*c)* ^ *Ccons (c))* ^ *C*_*re*_*f(c)*, corresponding to the individual anchor prediction, baseline cluster consensus, and localized refined prediction.

##### Bi-Consensus Mode

Defaulted if iterative refinement was bypassed. This required strict dual agreement:

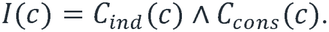

Finally, following identical compositional thresholding to remove sparse populations, the pristine counts matrix was subset to the global common ST gene intersection, transposed, and exported alongside dimensional metadata (sc_counts.csv, sc_labels.csv, st_counts.csv), providing a highly optimized, formatting-ready foundation for spatial deconvolution mapping.

### Integrated Variational Framework for Spatial Transcriptomic Deconvolution

#### Stage 1: Single-Cell Reference Encoding (scVAE)

##### Model Inputs

The scVAE receives three inputs from the reference scRNA-seq dataset. The expression matrix 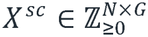 denotes the count matrix containing raw UMI counts for Acells across *G* genes. We deliberately model raw counts rather than normalized expression to properly account for count-based noise through the negative binomial likelihood. The cell type annotations *c ∈* {1,2,…, *C*]^*N*^ represent discrete cell type labels obtained through standard single-cell clustering, defining the identity that conditions the generative model. The library size 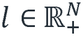 denotes the total UMI counts for each cell, computed as 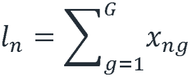. Explicitly modeling library size separates technical sequencing depth variation from biological expression differences.

##### Generative Model

We posit the following generative process for single-cell gene expression. For each cell rrwith cell type *c*_*n*_, the latent state 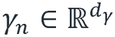 representing continuous cell state-capturing transcriptional variation within cell type *c*_*n*_ such as cell cycle phase, activation state, or metabolic status-is sampled from a standard normal prior:

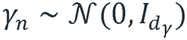

The expression rate decoding step computes gene expression rates through a deterministic mapping from latent space to the probability simplex:

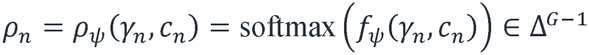

where 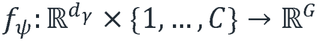 is a multi-layer perceptron mapping the latent state and cell type to unnormalized gene expression logits. The decoder architecture conditions on cell type through learned embeddings: *f*_*ψ*_(*γ, c*) = MLP_*ψ*_([ *γ*; *e*_c_]), where 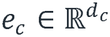 is a learnable embedding for cell type $c$.

Given the expression rate *ρ*_*n*_ and library size *l*_*n*_, the observed counts follow a negative binomial distribution parameterized by the mean *µ*_*ng*_ *= l*_*n*_ *· ρ*_*ng*_ and inverse dispersion *θ*_*g*_:

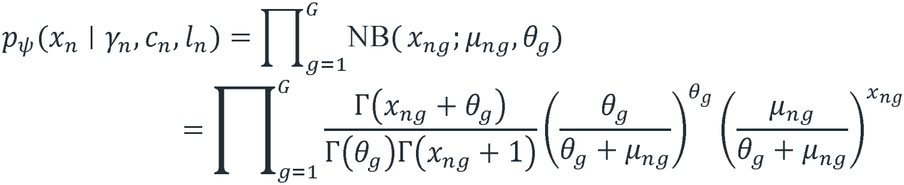

where *θ*_*g*_ *∈* ℝ_+_ is a learnable dispersion parameter (larger Vindicates less overdispersion).

Dispersion can be modeled as gene-specific (default), gene-and-cell-type-specific, or gene-and-batch-specific. Optionally, a zero-inflated negative binomial (ZINB) likelihood can be employed, modeling dropout probability through an additional neural network head.

##### Variational Inference

We employ amortized variational inference with an encoder network approximating the intractable posterior 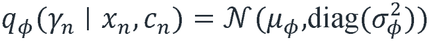. The encoder takes log-transformed counts, log (*x*_*n*_ + 1), and the cell type embedding *e*_*c*_ as inputs. We maximize the *β*-weighted evidence lower bound (ELBO):

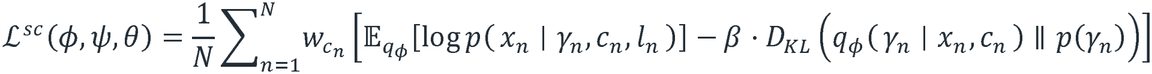

To handle class imbalance, we optionally apply inverse-frequency cell type weighting 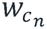 with a minimum proportion cap of 5%, ensuring rare cell types contribute meaningfully. To prevent posterior collapse, we employ KL annealing where the βweight is linearly increased from 0 to its target value over the first 50 epochs.

##### Empirical Prior Learning

After training the scVAE, we extract latent representations for all reference cells to fit cell-type-specific empirical priors. For each cell type c, we compute an empirical Gaussian prior 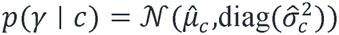 derived from the posterior means and variances of cells of type *c*(with an option to fit a Gaussian Mixture Model via Expectation-Maximization for complex distributions). These priors regularize inferred spatial states toward biologically plausible latent regions.

#### Stage 2: Warm-Start Initialization (stAE)

To stabilize the spatial deconvolution, we first compute reference expression profiles 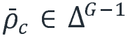 by averaging normalized counts across cells of each type. We then train a lightweight simplex autoencoder (stAE) that maps spatial expression directly to the probability simplex to estimate initial proportions:

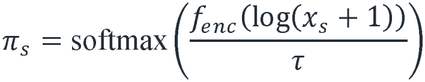

where τis a temperature parameter. The stAE minimizes a negative binomial NLL comparing the spatial spot to the library-size-scaled, α-corrected mixture of fixed profiles: 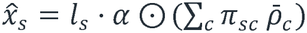, where *α ∈ ℝ*^*G*^ captures platform-specific gene correction factors.

#### Stage 3: Spatial Deconvolution and State Inference (stVAE)

##### Model Overview

The spatial VAE (stVAE) extends the framework to model mixed spatial spots. Each spatial spot sis modeled as a weighted mixture of cell types, where each cell type exhibits a distinct transcriptional state:

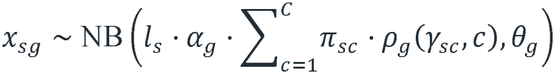

Gene correction factors *α*_g_ are constrained to [0.1,10.0]via clamping after exponentiation from log-space parameters, preventing numerical instability while allowing substantial platform-specific adjustments. Optionally, an “unknown” cell type category can be included to capture expression from cell types absent in the reference, initialized with a negative bias to discourage its use unless necessary.

##### State and Proportion Inference

Cell-type-specific latent states 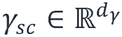 are inferred either through a shared amortized encoder network *f*_*gamma*_(log (*x*_*s*_ + l))or through direct non-amortized parameter optimization. Proportions *π*_*s*_ are initialized from the stAE warm-start estimates. Cell-type-specific expression profiles *ρ*_*sc*_ are then computed using the frozen scVAE decoder.

##### Loss Function and Constraints

The stVAE objective integrates reconstruction, prior regularization, and structural constraints:

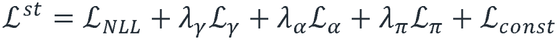

where *ℒ*_*NLL*_ the negative log-likelihood. The state penalty *ℒ* _*γ*_ *= Σ*_*s*_ *Σ*_*c*_ *π*_*sc*_ [−log *p(γ*_*sc*_ ∣ *c)]* regularizes states toward the empirical prior strictly for cell types present at the spot. ℒ_a_is an L2 penalty constraining log (*α*_*g*_ toward zero.

The proportion regularization *ℒ*_π_ enforces sharply resolved mixtures via two mechanisms. First, entropy minimization across cell types, computed as *H(π*_*s*_*) = − Σ*_*c*_ *π*_*sc*_ log *πs*_*C*_. Second, a Gini-coefficient-based sparsity penalty to explicitly encourage the absence of unrepresented cell types, computed as

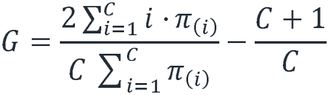

(where *π*_(*i)*_ are sorted proportions in ascending order). Maximizing Cencourages sparse mixtures.

To resolve the non-identifiability between changing cell states (*γ*) and cell proportions (*π*), we apply two-stage constraints (*ℒ*_*C*_*onst*). Users can impose hard constraints (freezing *π*_*s*_ and intensity profiles to stAE estimates) or soft constraints, which apply a Kullback-Leibler divergence penalty 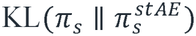.

##### Pseudo-Spot Consistency Training

To improve deconvolution accuracy, we incorporate consistency regularization using synthetic pseudo-spots. Pseudo-spots are created by randomly sampling 5-15 cells uniformly at random from the reference and summing their expression profiles 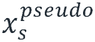, naturally reflecting cell type abundance. A supervised NLL loss *ℒ*_*pseudo*_ is evaluated on these spots against known ground-truth proportions 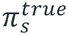 and added to the total loss via a progressively increasing warmup schedule.

### Downstream Analyses

#### Expression Imputation and Tissue Denoising

For each spot sand cell type c, the isolated expression profile is imputed as 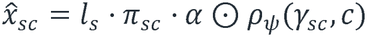 The aggregate expected counts *µ*_*sg*_ provide a denoised reconstruction of the tissue. Furthermore, cell-type intensity maps are computed as

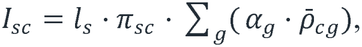

representing the expected total counts attributable to cell type *c* at spot s.

#### Visualization Suite and State Inference

The framework provides comprehensive downstream visualizations including: (1) spatial proportion maps with hexagonal tessellation matching Visium spot geometry, (2) cell type co-occurrence heatmaps, (3) UMAP/PCA projections of latent states *γ*_*sc*_, and (4) cell-type-specific spatial state gradients showing within-type transcriptional variation strictly within physical regions where the cell type resides.

#### Marker Gene and Differential Expression Analysis

To contextualize the inferred cell states, the pipeline conducts comprehensive reference-based marker gene ranking. Genes are scored integrating mean expression, specificity (ratio of expression in target vs. max of others), and a cell-type specific Gini coefficient. Statistical significance for differential expression is derived via one-vs-rest Wilcoxon rank-sum tests on the single-cell reference, adjusted using Benjamini-Hochberg FDR correction.

#### Implementation Details

Encoder and decoder networks utilize layer-normalized fully connected layers with ReLU activation and dropout (rate 0.1). Default architectures employ 2 hidden layers of 256 units, *d*_*γ*_ = 10, and *d*_*c*_ = 16. Numerical stability is ensured through epsilon constants (10^−8^), variance clamping (minimum 10^−4^), log-variance bounds [−10,10], and dispersion parameter clamping to [10^−4^, 10^4^].

Models are optimized via AdamW with gradient clipping (max norm 10.0). The stVAE uses decoupled learning rates based on a baseline 10−^3^rate with cosine annealing warm restarts (*T*_*o*_ = 100, *T*_*mult*_ = 2): *γ* networks (10^−3^), αand dispersion parameters (10^−4^), and proportions (5 x I0^−3^). Early stopping prevents overfitting by monitoring reconstruction loss with a patience of 30 epochs (scVAE) and 50 epochs (stVAE) and a minimum improvement threshold of 10^−4^.

Key hyperparameters include: proportion temperature *τ* = 0.5, *λ*_*π*_ = l0.0(entropy) and 1.0(Gini sparsity), *λ*_*γ*_ = 1.0, *λ* _*α*_= 0.01, and a pseudo-spot weight of l.0 phased over 50 warmup epochs.

### Spatial transcriptomics simulations and data generative process

#### Semi-synthetic ST generation

To rigorously evaluate the deconvolution fidelity of ARCADE against an absolute ground truth, we constructed a large-scale, semi-synthetic ST dataset. Because standard simulation approaches-which simply aggregate random subsets of single cells into “pseudo-bulks”-fail to capture the structured anatomical microenvironments and continuous transcriptomic gradients present in real tissue, we adapted the spatially-aware data generative framework originally introduced by *DestVI*^*17*^. The simulation utilizes the high-fidelity TCF scRNA-seq reference (generated in Stage 1) to populate a virtual spatial array, meticulously preserving both inter-cell-type distinctions and intra-cell-type continuous states.

#### Learning baseline profiles and continuous cell states

First, we extracted the data-generating parameters from the real TCF scRNA-seq reference. For every annotated cell type *c*, we calculated a baseline mean gene expression profile (*µc*) using counts-per-million (CPM) normalization. To capture the continuous transcriptomic variations (cellular states) that occur within a single cell type across space, we applied Principal Component Analysis (PCA) to the log-normalized counts of each cell type.^35^ This yielded a dense dictionary of within-cell-type transcriptomic variation (the PCA loadings, *W*_*c*_) and established the empirical bounds (mean and standard deviation) for the top five dimensions of cellular state embeddings.

#### Spatial topography and cell type proportions

To mimic the physical geometry of the 10x Genomics Visium platform, we constructed a staggered hexagonal lattice where the total number of capture spots operates as a tunable parameter (e.g., parameterized to 4,900 spots to represent a full Visium capture area). To simulate biologically realistic tissue architecture characterized by continuous spatial domains and gradual anatomical boundaries, we generated spatially autocorrelated cell type proportions using Gaussian Random Fields (GRFs).^36^ Specifically, for the spatial coordinates of each spot, we computed a squared-exponential covariance matrix to sample independent spatially smooth random fields for each cell type. These fields were interpreted as pre-normalized logits. To map these logits to valid cell type mixing proportions (*π*_*cs*_) for spot s, we applied a softmax function with a temperature parameter (*T* = 0.5):

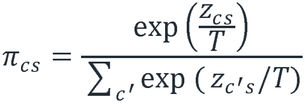

where *π*_*cs*_ represents the final ground-truth proportion (ranging from 0 to 1) of cell type cwithin spatial spot s. The variable z_cs_ denotes the unnormalized logit value derived from the Gaussian Random Field, representing the raw spatial intensity of cell type cat spot s. The temperature parameter T controls the sharpness of the spatial transitions; a value of *T =* 0.5was selected to enforce distinct dominant cell types while maintaining continuous gradients at region borders. Finally, the denominator iterates over c’, representing all available cell types in the reference, to normalize the proportions such that they sum to exactly 1 for every spot. This approach effectively eliminates the unrealistic “salt-and-pepper” noise common in naive simulations, resulting in contiguous, biologically plausible spatial domains. Identical spatial smoothing (GRFs) was applied to sample the continuous cell state embeddings (*γ*_*cs*_) for each cell type at every spot, ensuring that adjacent cells of the same type exhibit related transcriptomic states.

#### Synthetic expression and Negative Binomial count sampling

For each spatial spot s, we simulated a realistic cellular density by randomly drawing an absolute total of 5 to 20 cells. The specific quantity of each cell type within the spot was dictated by the ground-truth proportions (*π*_*cs*_). The target biological expression profile for a cell type *c* at spot *s* was computed by modulating the baseline expression profile (*µ*_*c*_) with the spatially generated cell state embeddings mapped through the PCA loadings 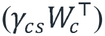.

Finally, to mimic the technical noise and sparsity inherent to spatial capture technologies, the target mixed expression for each spot was scaled by a capture efficiency factor of 10%.

Following established practices for modeling scRNA-seq and spatial transcriptomic count data [1, 5], the final observed discrete gene counts (*X*_S5_) for spot sand gene *g*were sampled from a Negative Binomial generalized linear model with a fixed overdispersion parameter:

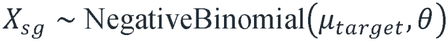

Here, X_S5_is the simulated integer read count for gene *g* at spatial spot *s*, representing the final synthetic data matrix. The mean parameter *µ*_*target*_ is the mathematically expected gene expression at that specific spot, calculated by combining the local cell-type proportions, baseline profiles, and spatial state modulations, scaled down by the capture efficiency. The overdispersion parameter *θ* (empirically set to 10 in our model) accounts for the high variance, technical noise, and characteristic dropouts (sparsity) typical of sequencing-based spatial transcriptomic technologies.

#### Computational implementation and parameterization

To enable comprehensive benchmarking across diverse spatial architectures, we developed the generative pipeline as a modular Python framework. This implementation allows for the manual parameterization of key experimental variables via command-line arguments, specifically the total spot count (--n_spots) and the cellular density per spot (--min_cells, --max_cells). For example, to simulate a standard 10x Visium array of the mouse striatum, the framework can be configured to generate 4,900 capture spots with a density of up to 10 cells per spot. By systematically tuning these arguments-adjusting spot counts to reflect varying tissue sizes, modifying density to test algorithmic resilience to cellular crowding, and utilizing divergent random seeds (e.g., seed 123) for statistical replication-we produced a diverse suite of paired scRNA-seq and ST matrices. Each simulation generates exact ground-truth values for spatial coordinates, cell-type proportions, and continuous cell states, providing a rigorous foundation for evaluating deconvolution fidelity across multi-scale scenarios.

#### Generation of Simulated scRNA-seq Data

To rigorously benchmark the ARCADE spatial deconvolution framework, a semi-simulation pipeline was developed to generate synthetic spatial transcriptomics (ST) data with known ground-truth parameters. The complete source code for this framework (simulation_ST_data.py) is available in the benchmarking/directory of the ARCADE GitHub repository. The simulation utilizes an authentic, user-provided single-cell RNA-seq (scRNA-seq) reference to compute baseline cell-type expression profiles and PCA-derived continuous cell state dimensions. To generate the synthetic spatial landscape, we simulated a staggered honeycomb lattice resembling the 10x Visium array geometry. Spatially smooth random fields were generated using a squared-exponential covariance function, which were then transformed to generate localized clusters of distinct cell types. Ground-truth spatial cell-type proportions were calculated via softmax normalization with temperature scaling, while continuous cell states were independently simulated using smoothed random fields projected onto the original scRNA-seq principal component space. Finally, the mixed spatial expression profiles were computed for each spot by linearly combining the simulated cell-type proportions and state-modulated expression profiles. Synthetic ST count matrices were then generated via negative binomial sampling to realistically model sequencing overdispersion and technical capture efficiency. This robust simulation framework outputs highly standardized spatial matrices-including raw intensities, normalized proportions, cell states, and SpaceRanger-compatible spatial coordinates-facilitating precise, objective evaluations of spatial deconvolution accuracy.

### Data analysis

#### 1. Read Alignment and Data Generation

For single-cell and single-nucleus RNA-seq analysis, processed reads were aligned using Cell Ranger (v9.0.1) and downstream analyses were conducted with Scanpy (v1.11.1). Simulated datasets were generated with Splatter (v1.26.0) and polyester (v1.38.0). For mouse datasets, reference genome resources were derived from the Ensembl Mus musculus GRCm39 assembly (including Mus_musculus.GRCm39.dna.primary.assembly.fa, Mus_musculus.GRCm39.cdna.all.fa, and Mus_musculus.GRCm39.112.chr.gtf) utilized within the refdata-gex-GRCm39-2024-A reference package. For human datasets, the refdata-gex-GRCh38-2024-A reference package was used, derived from the Homo sapiens GRCh38 assembly and GENCODE v44 annotation (specifically Homo_sapiens.GRCh38.dna.primary_assembly.fa.modified and gencode.v44.primary_assembly.annotation.gtf.filtered).

#### 2. scRNA-seq Data Quantification and Cell-Gene Matrix Generation

Sequencing reads were processed and quantified using the 10x Genomics Cell Ranger pipeline. The cellranger count command was executed to align reads to the respective reference transcriptomes, perform cell identification, and generate cell-gene expression matrices.

For the mouse brain dataset, the specific command used was:

~~~
cellranger count \
--id=5k_mouse_brain_CNIK_3pv3_output \
--sample=5k_mouse_brain_CNIK_3pv3 \
--fastqs=/home/data/qs/mus_str.sn/HGC20231021002-0003-2/cellranger_output/ \
--transcriptome=/home/data/qs/tools/refdata-gex-GRCm39-2024-A/ \
--localcores=40 \
--localmem=200 \
--expect-cells=5000 \
--create-bam=true
~~~

For the human PBMC dataset, the specific command used was:

~~~
cellranger count \
--id=pbmc_10k_v3_output \
--sample=pbmc_10k_v3 \
--fastqs=/home/data/qs/scRNA_seq_data_public/pbmc_10k_v3_fastqs/cellranger_output \
--transcriptome=/home/data/qs/tools/refdata-gex-GRCh38-2024-A \
--localcores=40 \
--localmem=200 \
--expect-cells=12000 \
--create-bam=true
~~~

#### 4. Python-Based Bayesian Optimization Framework

Post-quantification analysis and spatial reference generation were conducted using a unified, fully automated Python pipeline (01a_ARCADE_ref_optimizer.py) built upon the Scanpy and CellTypist frameworks. Replacing previously fragmented workflows, this integrated tool dynamically detects dataset structures, automatically applying Harmony integration for multi-sample or batched data while strictly enforcing global random seeds to ensure full computational reproducibility. The framework operates in two continuous stages. Stage 1 utilizes Bayesian Optimization (via scikit-optimize), executing simultaneous “Exploit,” “Explore,” and “BO-EI” search strategies to tune critical hyperparameters, including highly variable genes, principal components, nearest neighbors, and clustering resolution. The Bayesian objective function evaluates parameter fitness across configurable targets: a Biological Model maximizing the Cell Annotation Score (CAS) and Marker Confidence Score (MCS); a Structural Model incorporating Silhouette scores; and a novel MPS-Enhanced Model that utilizes an F1-based Marker Prior Score (MPS) combined with a composite DEG ranking scheme to reward the recovery of canonical marker genes. Stage 2 performs the final high-resolution analysis using the discovered optimal parameters and optionally triggers an iterative refinement loop, performing targeted sub-clustering on low-confidence (CAS-deficient) populations to resolve hidden or rare cell states. Ultimately, the framework employs a rigorous tri-consensus filtering strategy—retaining only cells exhibiting perfect agreement between individual machine-learning predictions, cluster-level consensus, and post-refinement labels. These ultra-high-confidence cells are seamlessly exported as standardized matrices (sc_counts.csv and sc_labels.csv), providing an optimized, noise-free reference for downstream spatial deconvolution.

#### 5. Seurat-Based Bayesian Optimization Framework (R Implementation)

To ensure broad accessibility within the R/Bioconductor ecosystem, we developed a parallel, Seurat-based (v5.0+) implementation of the ARCADE Reference Optimizer. This pipeline automates the tuning of critical scRNA-seq hyperparameters-including highly variable genes, principal components, nearest neighbors, and clustering resolution-via Bayesian Optimization, while dynamically applying Harmony integration to correct batch effects in multi-sample datasets. Unlike the Python implementation, this workflow leverages Seurat’s cross-dataset anchoring strategy to project reference cell-type labels, deriving robust cluster-level annotations through majority voting. To evaluate hyperparameter fitness, the Bayesian objective function maximizes the geometric mean of stability metrics by integrating three evaluative frameworks: a Biological Model driven by Cell Annotation and Marker Confidence Scores (CAS and MCS); a Structural Model utilizing Silhouette Scores to penalize poor manifold separation; and an MPS-Enhanced Model that evaluates canonical marker recovery using a novel Marker Prior Score weighted by log2 fold-change, expression level, and detection specificity. The engine employs simultaneous “Exploit,” “Explore,” and “BO-EI” search strategies to ensure robust convergence across the parameter space. Following optimal parameter discovery, an iterative refinement stage performs targeted sub-clustering on low-CAS clusters to resolve rare or poorly separated cell states. Ultimately, the pipeline applies a strict tri-consensus filtering strategy-retaining only cells with perfect agreement between individual predictions, cluster consensus, and post-refinement labels-to export highly confident, spatially-decoupled reference matrices for the Stage 2 ARCADE Spatial Decoupler.

#### 6. Spatial Deconvolution Data Export

To ensure transparency, reproducibility, and utility for downstream analysis, the outputs from the ARCADE Spatial Decoupler are comprehensively exported. Following the two-stage inference process (stAE and stVAE), the pipeline generates standardized CSV matrices containing cell-type proportion estimates, raw and normalized intensity maps, and spot-level cell state (y) continuous representations. Additionally, cell-type-specific imputed spatial expression matrices are exported alongside denoised tissue reconstructions. To facilitate biological interpretation, the pipeline automatically generates publication-ready spatial mapping plots, cell-type co-occurrence heatmaps, cell state continuum visualizations across UMAP embeddings, and top marker gene expression profiles.

### Collection of tissue samples

B6-hHTTCAG130 mice (Strain No. T054804, GemPharmatech), which harbor a human HTT gene fragment containing 130 CAG repeat expansions, were used as a Huntington’s disease model. Striatal tissues from these mice were collected and designated as musHuntingtonD-STR, while corresponding striatal samples from wild-type control mice were designated as musWT-STR. All animals were maintained under specific pathogen-free (SPF) conditions, provided with autoclaved food, and given acidified, autoclaved water. All experimental procedures involving animals were performed in strict accordance with institutional guidelines and approved by the Animal Care and Use Committee of Shenzhen University of Advanced Technology.

### snRNA-seq Library preparation

Single nuclei were isolated from the collected mouse striatum samples, and snRNA-seq libraries were prepared using the 10x Genomics Chromium Next GEM Single Cell 3’ v3.1 Kit according to the manufacturer’s protocol. In this workflow, individual nuclei were encapsulated in gel bead-in-emulsions (GEMs), lysed to release mRNA, reverse-transcribed into cDNA, and subsequently pre-amplified for downstream sequencing.

### Sequencing

The purified PCR libraries were sequenced on an Illumina NovaSeq 6000 platform using Paired-End 150 bp (PE150) chemistry. The sequencing was conducted to a predetermined data yield per library to ensure sufficient coverage for downstream analysis.

### Statistics

The statistical foundation of our study is the scBOA framework, which employs Bayesian Optimization (BO) to identify optimal hyperparameters (e.g., highly variable genes, principal components, neighborhood sizes, and clustering resolution) for scRNA-seq analysis. The BO process iteratively maximizes a multi-component objective function, f(x), which quantifies the biological and structural quality of the resulting cell partitions. This objective function integrates three key metrics: (1) the Cluster Annotation Score (CAS), a statistical measure of the enrichment, confidence, and specificity of known marker genes mapped to predicted cell types; (2) the Marker Prior Score (MPS), which assesses the fidelity of cluster markers against external canonical databases; and (3) the Silhouette score, which measures the structural density and separation of clusters in the latent space.

To rigorously quantify annotation fidelity, the MPS was formulated as an F1 Score-the harmonic mean of precision and recall. Precision was defined as the fraction of top differentially expressed genes (DEGs) that are recognized as canonical markers for the assigned cell type, while recall was defined as the fraction of known canonical markers successfully captured within the top DEGs. Following the identification of an optimal parameter set and final cell clustering, marker genes for each annotated cell type were identified using the non-parametric Wilcoxon rank-sum test (Mann-Whitney U test), as implemented in scanpy.tl.rank_genes_groups. This test evaluates the expression distribution of each gene within a given cluster against all other cells. The resulting p-values were corrected for multiple comparisons using the Benjamini-Hochberg procedure to control the false discovery rate (FDR), with an FDR < 0.05 considered statistically significant. To robustly rank these DEGs, a weighted composite statistical ranking method was utilized. Rather than relying solely on log-fold changes, this method normalizes and calculates a weighted linear combination of the log2 fold-change (40%), mean expression magnitude (30%), and cluster specificity (the percentage difference of cells expressing the gene, 30%).

Furthermore, to map predicted cell clusters to canonical marker databases, robust textual and semantic matching algorithms were employed. This included the calculation of the Jaccard similarity index for tokenized cell-type term overlaps, alongside Levenshtein distance-based fuzzy string matching (token set, token sort, and partial ratios) to ensure statistically robust cell-type synonym mapping. Where applicable, linear relationships between continuous variables were assessed using Pearson correlation coefficients, with adjusted R^2^ values used to evaluate model fit.

## Acknowledgments

This work was supported by funding from the National Key Research and Development Program of China (Grant Nos. 2023YFA0914904, 2022YFA1105601). Establishment of Ethical Framework and Policy System of Synthetic Biology (2024YFA0917200).

## Author contributions

Author contributions were as follows: Qiang Su conceived the project and designed the study. Qiang Su and Qizhou Lian supervised the study. Qiang Su performed the primary data analyses and generated the figures. Qiang Su and Yi Long wrote the initial draft of the manuscript, with subsequent input from all authors. Qiang Su, Jing Jiang and Chen Yang contributed specifically to the interpretation of the snRNA-seq data. All authors read and approved the final version of the manuscript.

## Data availability

The raw sequencing data generated for this study have been deposited in the NCBI Sequence Read Archive (SRA; https://www.ncbi.nlm.nih.gov/sra) under project accession PRJNA1263897. The snRNA-seq datasets for musHuntingtonD-STR and musWT-STR are available under this project with SRA accessions SRR33607933 and SRR33607934, respectively. Human prefrontal cortex data (specifically samples Br6432_anterior, Br6522_anterior, and Br2720_middle sample), originally described by Huuki-Myers et al. in A data-driven single-cell and spatial transcriptomic map of the human prefrontal cortex, were obtained from the National Institute of Mental Health (NIMH) Data Archive (https://nda.nih.gov) and the Globus endpoints “jhpce#spatialDLPFC” and “jhpce#DLPFC_snRNAseq” (accessible via http://research.libd.org/globus). Furthermore, datasets utilized for the concordance analysis of annotation fidelity (10x Genomics 5k Mouse Brain CNIK 3’v3.1 and 10k Human PBMC v3) are publicly available and were downloaded directly from the 10x Genomics Resource Dataset website.

## Code availability

The ARCADE computational pipeline is available as open-source software under the MIT License. The framework is implemented in Python and R, and can be executed in standard computational environments using the provided dependency files. Source code, comprehensive documentation, standardized benchmarking scripts, and example datasets are freely available on GitHub at https://github.com/QiangSu/ARCADE.

## Supplementary Materials

Figures S1-S46

Tables S1-S4

